# Coalescent-based branch length estimation improves dating of species trees

**DOI:** 10.1101/2025.02.25.640207

**Authors:** Yasamin Tabatabaee, Santiago Claramunt, Siavash Mirarab

## Abstract

Species trees need to be dated for many downstream applications. Typical molecular dating methods take a phylogenetic tree with branch lengths in substitution units as well as a set of calibrations as input and convert the branch lengths of the species tree to the unit of time while being consistent with the pre-specified calibrations. When dating species trees from multi-locus genome-scale datasets, the branch lengths and sometimes the topology of the species tree are estimated using concatenation. However, concatenation does not address gene tree heterogeneity across the genome. While Bayesian dating methods can address some forms of gene tree heterogeneity, such as incomplete lineage sorting, they are not scalable to large datasets. In this paper, we introduce a new scalable pipeline for dating species trees that addresses gene tree discordance for both topology and branch length estimation. The pipeline uses discordance-aware methods that account for incomplete lineage sorting for estimating the topology and branch lengths and maximum likelihood-based methods for the dating step. Our simulation study on datasets with gene tree discordance shows that this pipeline produces more accurate and less biased dates than pipelines that use concatenation or unpartitioned Bayesian methods. Furthermore, it is substantially more scalable and can handle datasets with thousands of species and genes. Our results on two biological datasets show that this new pipeline improves the inference of node ages and branch lengths for some nodes, in particular extant taxa, and improves the downstream diversification analysis.

## Introduction

Phylogenetic trees with branch lengths in units of time enable the study of evolutionary processes, such as species diversification, across geological times. The challenge (Forest, 2009) is to extrapolate from assumed dates for some nodes to the rest of the tree while modeling complex processes such as changes in substitution rates across lineages (Kumar, 2005; Langley and Fitch, 1974). A rich computational toolkit has been developed to deal with such complexities (Dos Reis et al., 2016; Ho and Duchêne, 2014; Kumar and Hedges, 2016; Rambaut and Bromham, 1998). Some of these methods take as input a tree with branch lengths measured in units of the expected number of substitutions per sequence site (SU), while others, typically Bayesian methods (Dos Reis et al., 2016), co-estimate the tree topology and branch lengths in units of time (Fig. 1A).

**Figure 1:**
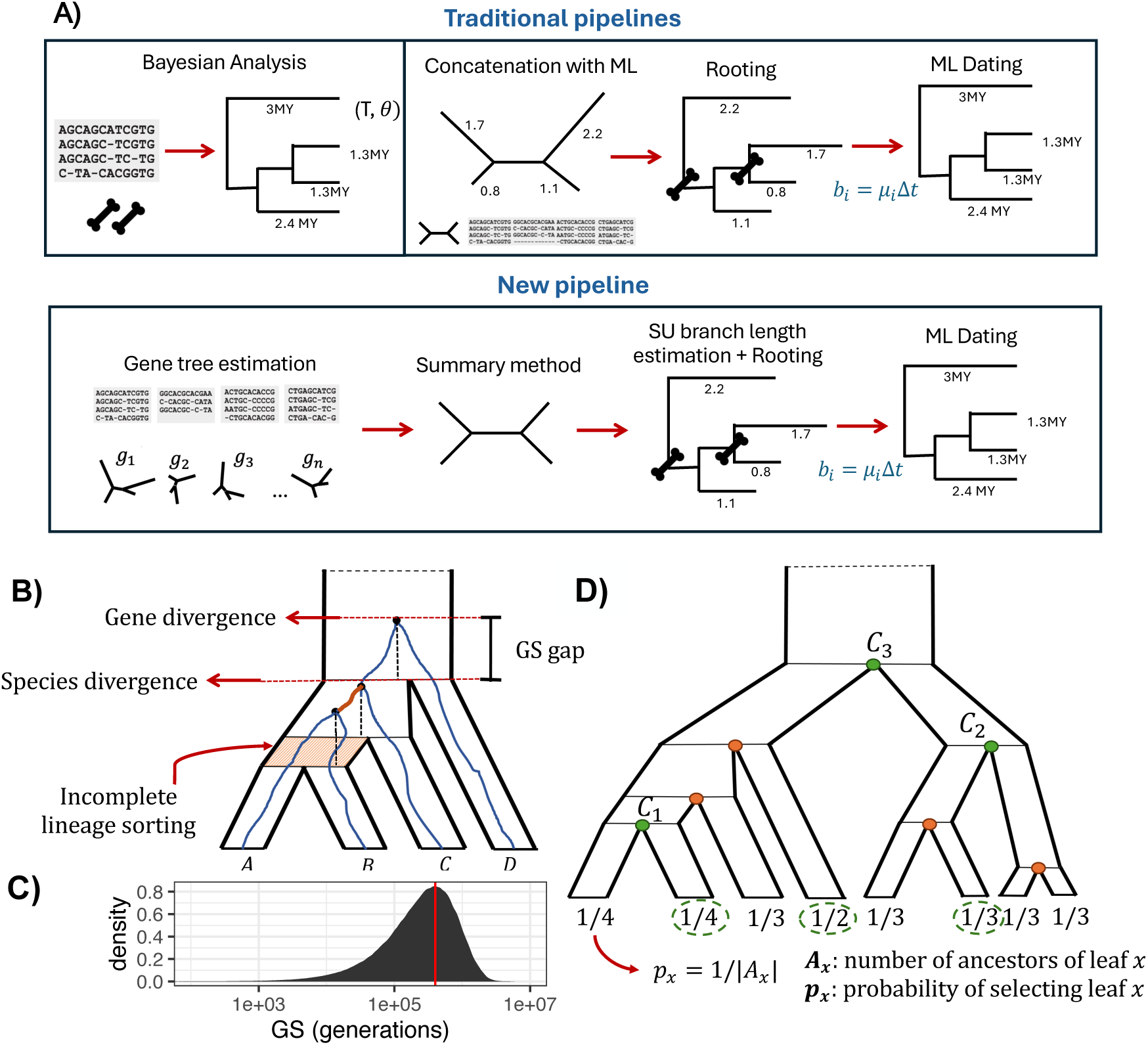
A) Traditional pipelines for estimating divergence times: Bayesian dating directly uses sequence data and calibration points and co-estimates the topology and branch lengths in time units; the two-step concatenation method estimates the branch lengths and/or topology using concatenation with maximum likelihood to produce a tree with branch lengths in units of number of substitutions per site, and then dates this tree by transforming these branch lengths. The scalable four-step pipeline introduced in this study uses discordance-aware methods for both topology and branch length estimation. B) Gene tree evolving inside a four-taxon species tree and the gap between gene divergence and species divergence (GS-gap). While incomplete lineage sorting can increase the GS-gap, this gap exists even when there is no ILS. Due to the GS-gap, terminal branches in the gene trees are always longer than their corresponding branch in the species tree, but internal branches may be shorter or longer (e.g., the internal branch colored in orange is shorter in the gene tree). C) The distribution of GS gap measured in units of generations is shown for terminal branches of 101 species, 50 replicates, with 1000 gene trees per replicate; *N_e_* = 4 *×* 10^5^ (red line). D) Protocol for simulating calibration points on a species tree topology. Each leaf *x* is assigned a probability *p_x_* = 1*/|A_x_|* where *A_x_* is the number of nodes between *x* and the root (inclusive). Then, *k* taxa are selected randomly based on the probabilities assigned to them (in the figure, *k* = 3 and the three taxa with green circles are selected). Then for each selected taxa *x*, one of its *A_x_* ancestors is selected with probability 1*/A_x_* (the green internal nodes). With this process, *k* internal nodes are selected and the true divergence times of these nodes are used as calibration points.

A challenge facing accurate dating is the impact of the heterogeneity of evolution across the genome. Beyond well-documented substitution rate differences across the genome (Rasmussen and Kellis, 2007; Thorne and Kishino, 2002; Wolfe et al., 1989), different regions can have different evolutionary histories (Maddison, 1997). In particular, each recombination-free region of the genome (a locus) has its own coalescence history, potentially including incomplete lineage sorting (ILS) and topological differences across the genome. This stochastic process can be studied using the multi-species coalescent (MSC) model (Rannala and Yang, 2003). Under this model that considers coalescence *times* and not just topologies, each locus has a *unique* coalescent history; although they can be arbitrarily close, no two loci have exactly the same history.

It has long been appreciated that there is a gap between gene divergence and species divergence times (Arbogast et al., 2002; Edwards and Beerli, 2000) (Fig. 1B). Gene genealogies from any two species will coalesce some time *earlier* than the corresponding speciation event. Thus, gene divergences always predate species divergences, and this gene/species (GS) gap is in expectation proportional to the ancestral effective population size (Fig. 1C); the expected length of the GS gap under the Wright-Fisher model for two lineages is a single coalescent unit (CU). Note that the GS gap does not impact all branches equally due to several reasons, including varying population sizes. More significantly, the terminal branches in the gene tree are always longer than the terminal branches in the species tree. In contrast, internal branches in the gene tree may or may not be present in the species tree, and when present, they may be longer or shorter (Fig. 1B). Thus, ignoring the GS gap will not just add random noise but rather will bias the dates: Terminal branch lengths are expected to be systematically over-estimated.

Modeling the impact of heterogeneity due to ILS has been an active area of research, leading to many methods to infer species trees (Mirarab et al., 2021; Rannala et al., 2020). These methods broadly fall into three categories: Bayesian methods that co-infer species trees and gene trees with parameters corresponding to divergence times and population sizes, two-step methods that first infer gene trees and then summarize them to obtain a species tree, and site-based methods that infer a species tree without inferring gene trees. While all these methods infer the species tree topology and are statistically consistent estimators, many have paid scarce attention to branch lengths. Concatenating loci as one input matrix, in contrast, is a statistically inconsistent estimator of topology (Roch and Steel, 2015); it produces branch lengths but suffers from the GS gap bias mentioned earlier. We can expect that using concatenation for dating will introduce systematic biases, including over-estimation of terminal branch lengths.

This bias in dating terminal branches is not just a theoretical concern and has been observed empirically (Ho et al., 2005; Ksepka et al., 2014; Mendes and Hahn, 2016; Phillips, 2009; Schwartz and Mueller, 2010). It can impact downstream analyses, such as the estimation of lineage diversification dynamics. In particular, the GS gap may contribute to the apparent pattern of early burst and slow-downs in diversification shown by many empirical phylogenies based on gene trees, as the long terminal branches result in lower estimates of speciation rates (Burbrink and Pyron, 2011). In addition, the overestimation of terminal branches may be behind the inability to detect extinction from molecular phylogenies, because extinction manifests itself as an excess of short terminal branches in reconstructed phylogenies (Nee et al., 1994).

Bayesian methods for inferring species trees under MSC (Flouri et al., 2018; Ogilvie et al., 2017; Rannala and Yang, 2003) have always modeled branch lengths using explicit parameters and thus can produce correct species divergence times. In fact, inferring correct branch lengths has been one of their main advantages (Ogilvie et al., 2017) over two-step approaches. Their main shortcoming is that they are far less scalable than the two-step approach, which has scaled to thousands of species (e.g., Harvey et al., 2020; Zhu et al., 2019; Zuntini et al., 2024) and tens of thousands of loci (e.g., Stiller et al., 2024). However, because gene tree summarization methods have not traditionally produced reliable branch lengths, users have needed to resort to concatenation for inferring branch lengths in substitution units (e.g., Song et al., 2012; Stiller et al., 2024) for dating. Using concatenation to get branch lengths makes the two-step approach prone to the negative effects of the GS gap. Finally, some likelihood-based methods (Peng et al., 2022; Schrempf et al., 2016) allow site-based methods to infer divergence times and are more scalable than Bayesian methods; yet, these are not as scalable as the existing two-step methods. Therefore, existing scalable approaches have not provided unbiased branch lengths and dates and approaches that do provide unbiased branch lengths are not scalable.

Recent advances hold the promise of enabling *scalable and unbiased* branch length estimation in both substitution and time units. In particular, our new method, CASTLES-Pro, can compute SU branch lengths by summarizing gene tree branches while accounting for the GS gap (Tabatabaee et al., 2025). Simulation studies showed that it is far more accurate than concatenation (and similar methods) in the presence of sufficiently high ILS (Arasti et al., 2024). It reduced terminal branch lengths systematically compared to concatenation, as one would expect. At the same time, several scalable methods of dating exist, including those based on maximum likelihood (ML) (e.g., Mai et al., 2024; Sagulenko et al., 2018; Smith and O’Meara, 2012; Volz and Frost, 2017). Combining these advances, scalable branch length estimation and ML-based dating, promises a scalable and accurate approach for dating species trees that accounts for the GS gap.

In this paper, we study a scalable four-step approach for producing dated phylogenomic trees relying on new methods for SU branch length estimation accounting for coalescence (Fig. 1A). The accuracy and scalability of this pipeline have not been studied before. We show in extensive simulations that with enough ILS, this approach produces divergence times that are more accurate than using concatenation for branch lengths, as is commonly done. In addition, our proposed pipeline is far more scalable than pipelines based on concatenation, easily scaling to datasets with 10,000 species and 1000 genes when paired with fast ML-based dating methods. On real data, we show that the approach leads to substantially shorter terminal branches and eliminates artifactual inferences of sudden drops in speciation rates close to the present time.

## Material and Methods

Our proposed four-step pipeline using coalescent-based branch lengths (CoalBL) is as follows:

**Step 1.** Infer gene trees for individual loci (e.g., using RAxML or IQ-TREE).

**Step 2.** Summarize the gene trees to get the species tree topology (e.g., using ASTRAL).

**Step 3.** Estimate the expected number of substitutions per site along the branches of the tree while accounting for the GS gap (e.g., using CASTLES-Pro).

**Step 4.** Date the tree using a scalable ML-based method (e.g., TreePL, LSD, MD-Cat, or wLogDate).

We used both simulated and biological datasets to compare the proposed four-step dating pipeline against the standard pipeline that uses concatenation for branch length estimation (ConBL). Note that the ConBL approach can infer the species tree topology using either concatenation or summary methods. Thus, it can share steps 1 and 2 with our proposed pipeline but uses concatenation for step 3; alternatively, it can replace steps 1–3 with a single ML analysis using concatenation. Both pipelines share step 4.

### Study design: pipeline variations

We tested several variations of the proposed pipelines. We ensured ConBL and CoalBL are compared on the same underlying topology. For simulated datasets, we dated the true species tree topology for both pipelines, thus eliminating step 2. On biological datasets, we performed dating on topologies estimated using ASTRAL or concatenation. The species trees are rooted before dating, using either the true rooting (in simulations) or outgroups (for biological datasets). For both simulated and biological data, we used estimated gene trees available from prior studies, therefore eliminating the need to perform step 1.

The main difference between the two pipelines is the SU branch length calculation (step 3). For step 3 of CoalBL, we used gene trees as input to CASTLES-Pro, which is a scalable method that explicitly models the GS bias. For ConBL, we used RAxML (Stamatakis, 2014) applied on a fixed tree topology to estimate SU branch lengths. We also tested the Bayesian molecular dating using MCMCtree (Yang and Rannala, 2006) on a subset of our datasets. The input to MCMCtree is the tree topology (without branch lengths) and the sequence data; it bypasses step 3 and directly uses sequence data to date the topology. Unlike the ML-based methods that use fixed calibration points or min-max bounds, MCMCTree uses soft fossil calibrations with flexible probability distributions to describe the uncertainty in fossil ages. We used MCMCTree with approximate likelihood calculation (Reis and Yang, 2011; Thorne et al., 1998) to speed up the analyses (suggested for large inputs). All parameters of the MCMCTree analyses in simulations were taken from Stiller et al. (2024) study, with the exception of burnin, sample frequency, and the number of samples in the MCMC sampling that were set to 50,000, 100, and 50,000, respectively.

For dating (step 4), we examined four ML dating methods: LSD (To et al., 2016), which can be considered ML under a strict clock with Gaussian estimation noise (run in the QPD* mode and minimum branch length set to 0.001); wLogDate (Mai and Mirarab, 2021), which is ML under a LogNormal rate model; MD-Cat (Mai et al., 2024), which is ML under a categorical model of rates with *k* (default 50) rates; and TreePL (Smith and O’Meara, 2012), which uses a likelihood framework with penalties for rate divergences (Sanderson, 2002). These methods take as input a tree topology with branch lengths in substitution units and (optionally) a set of calibration points. For dating methods that need sequence length as input (LSD and TreePL), we specified the sequence length as the total number of sites across all genes.

In total, we compared nine dating pipelines (four ML-based dating methods with CoalBL and ConBL, as well as MCMCtree). For both ConBL and MCMCtree, we used unpartitioned analysis, where all gene sequences were concatenated and passed as a single partition to the method. Further details about each method and how we ran them are provided in Supplementary Appendix Section S1.1.

### Simulated datasets

We examined three simulated datasets with gene tree discordance due to ILS that were generated using the simulation software SimPhy (Mallo et al., 2016). These datasets have model conditions that vary in terms of the level of ILS, deviation from the clock, number of species and genes, and the amount of gene tree estimation error (GTEE). To evaluate the accuracy of dating, we need to have ground-truth species tree topologies in the unit of time. SimPhy generates true species trees in the unit of the number of generations and true gene trees in substitution units. To convert the species trees to units of million years, we multiplied the generation unit lengths by 5 (assumed generation time) and divide them by 10^6^. We created model conditions with different numbers of calibration points, ranging from zero to 3, 5, and 10 calibration points for small datasets and up to 50 calibrations for large datasets. When the ML-based dating methods are used without any input calibration points, they produce a unit-length ultrametric tree (MCMTree needs calibrations). While some dating methods can use calibrations in the form of min-max bounds or probability distributions, we specified the calibration points as fixed values everywhere.

To select *k* calibration points on the species tree, we first assigned a probability to each terminal node (leaf) inversely proportional to the number of nodes from the root to that leaf; we then randomly sampled *k* leaves based on these probabilities. For each selected leaf, we picked one node along the subtending lineage uniformly at random but avoided the ones already selected, if any (Fig. 1D). This procedure helps distribute the calibration nodes widely across the tree; if sampled uniformly at random, most calibration nodes would be placed near the tips and in species-rich clades. We then used the true divergence times of the sampled nodes as calibration points. In some of the experiments, we fixed one calibration point on the root of the tree and selected the *k −* 1 other calibration points using the approach described above. We refer to these experiments as “root-fixed”, as opposed to “root-unfixed”. For the root-unfixed experiments, we set a soft upper bound for the root age for MCMCTree as 10 + *T* where *T* is the true root age. Unless otherwise noted, results are for the *root-fixed* experiments.

We measure the error of dating using mean absolute error, bias, RMSE, and mean absolute logarithmic error, averaged across all species tree branches compared to the true tree. Absolute error and bias emphasize longer branches more than short branches, and RMSE and log error emphasize shorter branches more. Since the higher levels of ILS correlate with shorter tree heights, we report branch length error normalized by the species tree height (including the outgroup edge, when present). In addition to branch length error, we report errors in node ages, using the same four metrics. We also report the time of the most recent common ancestor (tMRCA) for the *root-unfixed* experiments. Finally, we report *treeness* (i.e., sum of internal branch lengths divided by sum of all branch lengths) of the dated trees. While neither CASTLES-Pro nor ConBL produce zero or negative SU branch lengths, some of the dating methods (in particular, LSD) can produce zero-length branches in time units. For all methods, we replace negative and zero lengths with the small pseudo-count of 10*^−^*^6^ before calculating error metrics.

#### 30-taxon dataset

We used a 30-taxon dataset from Mai et al. (2017) with six model conditions, varying in terms of deviation from the clock and inclusion of an outgroup. The gene trees in this dataset were estimated using FastTree-2 (Price et al., 2010) from alignments with gene sequence lengths that were drawn from a log-normal distribution (average: 495bp). The ILS level, measured in terms of Robinson and Foulds (1981) (RF) distance between the true species tree and true gene trees (denoted average distance or AD) was heterogeneous across replicates, with a mean of 46%. The average GTEE level is 38% across all model conditions and replicates. This dataset has three levels of deviation from the clock, specified with parameter *α* of the gamma distribution for branch rate heterogeneity, set to 5 (low), 1.5 (medium), or 0.15 (high). The number of genes in this dataset is 500 and the number of repliclates is 100. For this dataset, we created conditions with 0, 3, or 5 calibration points, both *root-fixed* and *root-unfixed*. The total number of model conditions for this dataset is 30 (2 outgroup settings *×* 3 *α* values *×* 5 calibration settings).

#### 100-taxon dataset

We used the 101-taxon (100 ingroup and 1 outgroup) dataset from Zhang et al. (2018) (50 replicates) with model conditions that vary in terms of the level of gene tree estimation error (GTEE). The four model conditions have GTEE levels of 23%, 31%, 42%, and 55%, respectively, corresponding to gene trees estimated using FastTree-2 from sequence alignments of length 1600bp, 800bp, 400bp, and 200bp, respectively. The level of ILS varied between 30% to 58% across replicates (mean: 46%). We used 0, 3, or 10 calibration points both as *root-fixed* and *root-unfixed* (for 3 and 10). The default number of genes in this dataset was 1000, which we also subsampled to get 500, 200, and 50 genes for the *root-fixed* experiments. In total, we created 56 different model conditions for this dataset.

#### Large dataset

To evaluate scalability, we generated a new dataset with large trees with up to 10,000 species using a modified version of the simulation software SimPhy created by Tabatabaee et al. (2023) that creates species trees with SU and CU branch lengths. Species trees are generated using a birth-death process, with birth and death rates sampled from log-uniform distributions. To simulate heterogenous levels of ILS, we draw the effective population size from a uniform distribution between 20,000 and 2,000,000. We simulated conditions with 50, 100, 200, 500, 1K, 2K, 5K, and 10K ingroups and 1 outgroup. We simulated 500bp gene sequence alignments using INDELible (Fletcher and Yang, 2009) under the GTR+Γ model with model parameters estimated from three different real datasets (identical to parameters used in Mirarab and Warnow (2015)) and then used FastTree-2 to estimate gene trees under the GTR model. The ILS level is highly heterogeneous across replicates (minimum: 0%, maximum: 92.35%), with an average of 47.26%, and the GTEE level is, on average, 46.47% across all model conditions. For a model condition with *n* taxa, we simulated 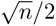 calibration points, resulting in 3, 5, 7, 11, 15, 22, 35, and 50 calibrations for the eight tree sizes, respectively. This dataset has 20 replicates for each model condition and 1000 genes in each replicate. Tables S1 and S2 summarize further information about this dataset. We ran all methods given 64GB of memory across 16 cores. For this scalability experiment, we only used TreePL, which was one of the fastest and most accurate dating methods.

### Biological datasets

We examine two avian biological datasets representing rapid radiations at two different timescales. We studied a dataset of modern birds (Neornithes) by Stiller et al. (2024), including 363 bird species and 63,430 genes. The main species tree used in the original study was estimated using ASTRAL and furnished with SU branch lengths using RAxML run on the concatenation of all 63K genes. Dating analyses in the original study were performed using MCMCtree with approximate likelihood calculation and a sequential-subtree approach using 34 calibration points. The tree was divided into 11 subtrees (with 19–42 taxa in each), and an initial dating analysis was performed on a backbone tree with 56 taxa containing two representatives of each of the 11 subtrees. The subtrees were then attached to the backbone tree to get a time tree on all 363 taxa. The analyses were done using a subset of the 10,494 most clock-like loci (i.e., those with the lowest coefficient of variation in root-to-tip distances).

We furnished the main ASTRAL topology from the original study with CASTLES-Pro branch lengths (using all 63K loci) and used three ML-based dating methods, MD-Cat, TreePL and wLogDate to estimate divergence times. While MCMCtree can take calibration densities in the form of probability distributions as input, the ML-based methods either work with fixed calibration bounds (MD-Cat and wLogDate) or minmax bounds (TreePL). Due to this difference, we cannot directly compare the ML-based trees to the output of MCMCtree, and mainly focus on comparing the dated trees based on ConBL or CoalBL branch lengths. To generate fixed calibration points and minimum-maximum bounds for the ML-based dating methods, we used the 0%, 50%, and 95% quantiles of the probability distributions of each fossil calibration density used in the original MCMCtree analysis. We used the median values (50% quantiles) as calibrations for MD-Cat and wLogDate that use fixed calibration points, and both median and min-max values (0%-95% quantiles) for TreePL that use min-max bounds. We examine variations in the age of orders, families, and genera, the correlation between terminal and internal branches, and changes in substitution rates across time.

We also study a suboscines dataset of Harvey et al. (2020), including 1940 individuals from 1283 species (98.2% of all suboscines species, which present the largest bird radiation in the tropics) with 2389 orthologous loci. Harvey et al. (2020) used this dataset to study the diversification of suboscines through time, and found that diversification rates (including speciation and extinction rates) have been relatively constant across the history of the group (between 40 to 51 million years) except for a dramatic drop in the past 2 million years. They suggested that this drop could have resulted from incomplete sampling or unsorted ancestral polymorphism. They also attribute this drop to challenges in studying diversification histories in extant taxa (terminal branches). Here, we re-analyze this dataset to see if this drop can be an artifact of a potential overestimation bias in branch lengths estimated using ConBL and whether CASTLES-Pro can reduce or eliminate this problem.

The original study had used both concatenation (with EXaML) and ASTRAL to estimate a species tree topology, but since the concatenation topology better matched previously known relationships, the concatenation tree was used for the final diversification analyses. The species trees were dated using TreePL, with four calibration points. We used CASTLES-Pro to estimate branch lengths on the ASTRAL and concatenation trees and used ConBL as provided by the original paper for both trees. We then used TreePL with the same calibration points, smoothing parameters and cross-validation method as the original study to date these two species trees. We then repeated the diversification-through-time analyses performed in Harvey et al. (2020) using a Bayesian MCMC analysis on the different species tree topologies furnished with ConBL or CoalBL branch lengths (in total four trees). We study the trend in the speciation rates and extinction rates by performing CoMet analysis using the TESS library (Höhna et al., 2016) where the diversification rates are estimated using reversible-jump MCMC. We remove the outgroup before the CoMet analysis and use automatic empirical hyperprior search and set the maximum number of iterations to 1,000,000. We also examine differences between terminal and internal branch lengths as well as lineage-through-time plots.

## Results

### Simulation studies

#### 30-taxon Dataset

On this dataset, using the CoalBL pipeline instead of ConBL decreases the mean absolute error of branch lengths in all conditions when the level of ILS is at least moderate (Fig. 2 and Fig. S1). As the level of ILS increases, the height-normalized error for all methods increases, but more slowly for CoalBL pipelines. For the lowest level of ILS (*<* 0.25 AD), ConBL pipelines have a small advantage, whereas, for remaining ILS levels, pipelines that use CoalBL are always advantageous, regardless of the dating method. As expected, more calibration points lead to lower errors for all methods; however, with high ILS, CoalBL with fewer calibrations is better than ConBL with more. Increased deviations from a strict molecular clock increase the error for all methods, and the positive impact of CoalBL is more substantial for conditions with low deviation from the clock (Fig. 2a).

**Figure 2:**
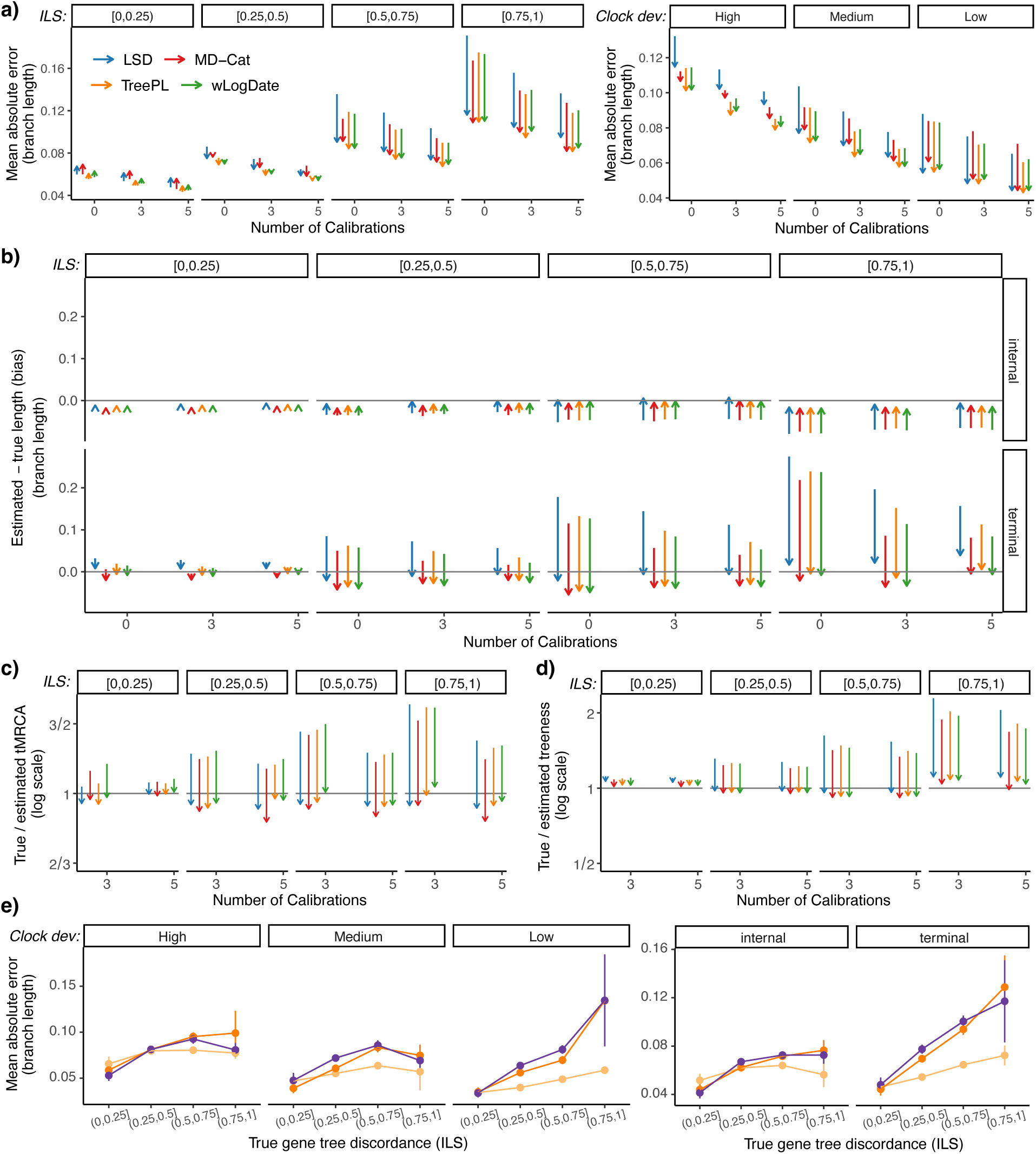
Results on 30-taxon simulated ILS datasets for conditions without outgroup. a) Change in height-normalized mean absolute error of branch lengths in the dating pipelines going from ConBL (arrow tail) to CoalBL (arrow head). Conditions vary in terms of the level of ILS (left: boxes), deviation from the clock (right: boxes), and number of calibrations (*x*-axis). b) Change in bias of branch lengths from ConBL to CoalBL. c) Change in true tMRCA (time of the most recent common ancestor) divided by estimated tMRCA from ConBL to CoalBL for the *root-unfixed* experiments in log2 scale. d) Change in true treeness (sum of internal branch lengths divided by sum of all branch lengths) divided by estimated treeness from ConBL to CoalBL for the *root-fixed* experiments in log2 scale. e) Comparison between mean absolute error of MCMCtree, TreePL+CASTLES-Pro and TreePL+ConBL for different levels of ILS, deviation from the clock, and branch type for experiments with five calibration points. Panels a, b, d, and e are for *root-fixed* experiments, while c is for *root-unfixed* (since tMRCA is fixed for *root-fixed*). The number of genes is 500 (100 replicates). See also Figs. S1-S7 for additional error metrics and other ways of dividing the data.

The reduction in error is mostly due to better terminal branch lengths (Fig. S2). The internal branch lengths change less and improve only in the two conditions with the highest ILS and the condition with the lowest deviation from the clock. Using ConBL leads to an overestimation bias for terminal branches and a smaller underestimation bias for internal branches, and this bias increases as the level of ILS increases (Fig. 2b). Because of these biases, treeness is underestimated in all pipelines based on ConBL (Fig. 2d) and for MCMCtree (Fig. S3), especially for high levels of ILS. The use of CoalBL reduces the underestimation bias of internal branches and eliminates the overestimation for terminal ones, often causing a relatively small underestimation bias for terminal branches (Fig. 2b). The CoalBL pipeline leads to relatively accurate levels of treeness, with a small overestimation bias for high deviation from the clock or ILS (Fig. S3). Deviation from the clock has very little to no impact on the treeness when using ConBL or MCMCtree; CoalBL pipelines change from no bias for conditions with low clock deviation to a small overestimation bias in conditions with high deviation.

Similar trends in bias are observed with or without an outgroup when the root is fixed in terms of treeness (Fig. S3) and bias (Fig. S4). However, not fixing the age of the root changes the patterns to some degree. In the *root-unfixed* conditions, the ConBL pipeline has a much larger underestimation bias for internal branches but lower levels of overestimation bias for terminal branches.

The inferred tMRCA (meaningful only for *root-unfixed* experiments) is substantially underestimated in all pipelines based on ConBL and slightly overestimated in most CoalBL pipelines (Fig. 2c). MCMCtree also substantially underestimates tMRCA (Fig. S5). The error in tMRCA becomes larger for all methods, especially those based on ConBL, as the level of ILS increases. Increasing the number of calibrations and decreasing deviation from the clock reduces the error of tMRCA for all methods, regardless of whether an outgroup is included (Fig. S5).

The relative accuracy of dating methods is variable and depends on the number of calibrations and level of ILS (Fig. 2a). Overall, the strict-clock LSD method is clearly the least accurate method, while TreePL is the most accurate, followed by MD-Cat. Moreover, patterns of error remain largely similar for the RMSE error and mean log error that emphasize shorter branches more than absolute error (Fig. S1). For the log error, LSD has very high errors (perhaps due to the abundance of near-zero lengths), and pairing it with ConBL or CoalBL makes very little difference. The trends also remain similar if we examine node ages instead of branch lengths (Fig. S6), focusing on absolute error and RMSE. There is, however, a somewhat larger underestimation bias for CoalBL pipelines when evaluating node age.

We compare MCMCtree with the most accurate ML-based dating method in our simulations, TreePL (Figs. 2 and S7). The comparison between MCMCtree and TreePL+ConBL depends on the deviation from the clock, with TreePL+ConBL performing better in more conditions overall. TreePL+CoalBL is more accurate than the other two methods for all levels of deviation from the clock and ILS, except for the lowest level of ILS (*<* 0.25 AD). These trends are consistent across different numbers of calibrations and when assessing node ages (Fig. S7). The bias of MCMCtree and ConBL are very close in all conditions, with MCMCtree performing slightly better for internal branches and ConBL for terminal ones (Fig. S7). Biases of both internal and terminal branches are substantially reduced when using TreePL+CoalBL.

#### S100

On the 100-taxon datasets, with 1000 genes, using CoalBL instead of ConBL reduces the mean absolute error of branch lengths and node ages across most sequence length levels and dating methods (Fig. 3a). Increasing the number of calibration points reduces the error for all dating pipelines, and the advantage of CoalBL is larger when using fewer calibrations. Given poorly estimated gene trees from 200bp genes and 10 calibration points, CoalBL and ConBL are tied. As the sequence length increases and GTEE decreases, dating using CoalBL improves far more than ConBL, such that with 1600bp, CoalBL has a clear advantage even with 10 calibration points. The trends for RMSE and log error are similar, though the magnitude of improvements differs in each case (Fig. S8).

**Figure 3:**
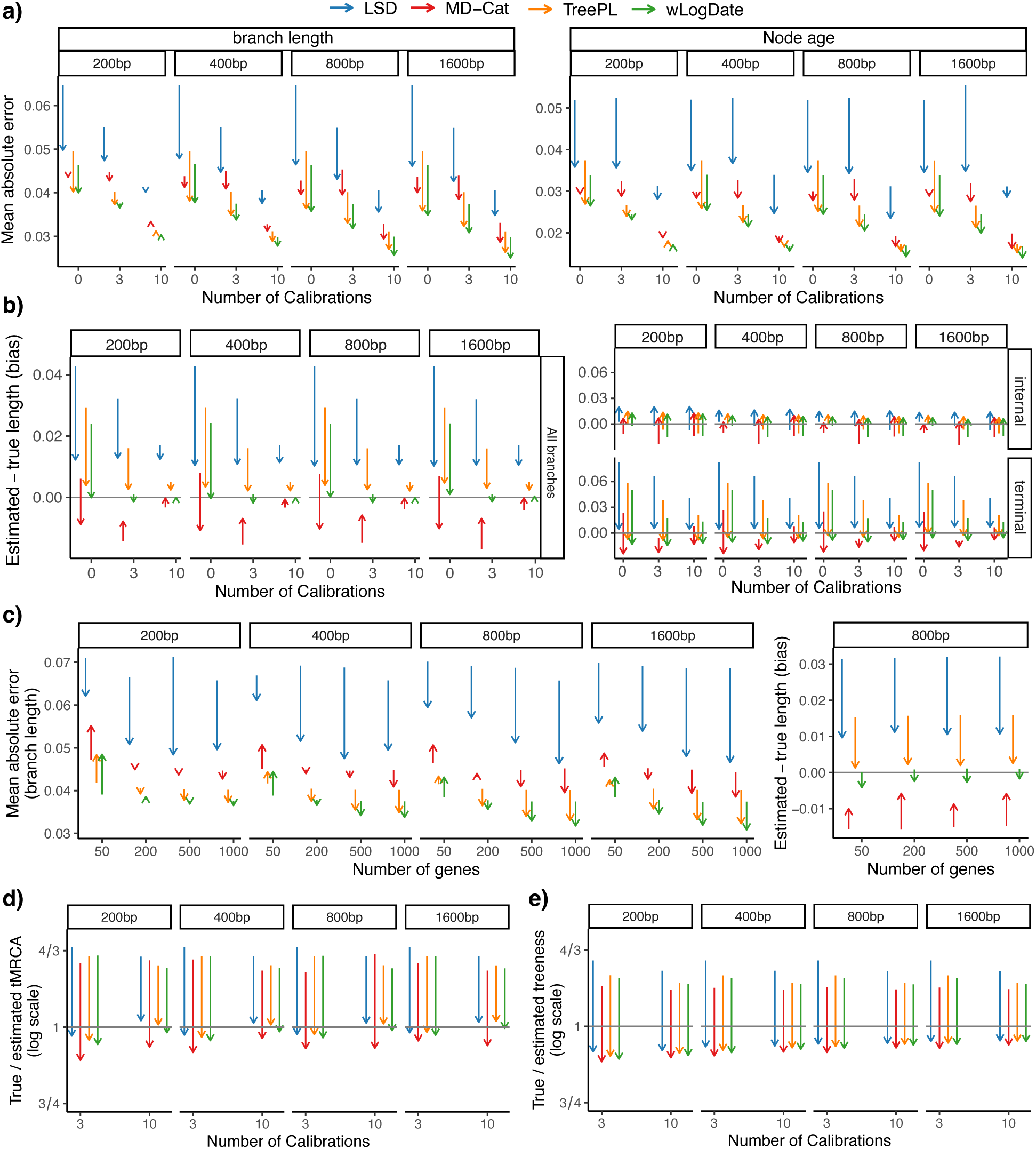
Results on 100-taxon ILS simulated datasets. a) Change in height-normalized mean absolute error of branch lengths and node ages in the dating pipelines when going from ConBL (arrow tail) to CoalBL (arrowhead) in the root-fixed experiments. Conditions vary in terms of the sequence length and number of calibrations. The number of genes is 1000. b) Change in bias of branch lengths for conditions with different sequence lengths in the *root-fixed* experiments. The number of genes is 1000. c) Change in mean absolute error and overall bias of branch lengths for conditions with different numbers of genes and sequence lengths in the *root-fixed* experiments. The number of calibrations is 3. d) Change in tMRCA ratio (true tMRCA divided by estimated tMRCA) for the *root-unfixed* experiments. e) Change in treeness ratio (true treeness divided by estimated treeness) for the *root-fixed* experiments. Conditions vary in terms of sequence length and number of calibrations. See also Figs. S8-S10 for more data.

In terms of bias, using CoalBL instead of ConBL reduces the overall bias for all dating methods in all conditions when calibration points are used (Fig. 3b), whether this bias is towards overestimation or underestimation. With the exception of MD-Cat, pipelines based on ConBL have a small underestimation bias for internal branch lengths and a larger overestimation bias for terminal branches. Both biases are substantially reduced with CoalBL and are either eliminated or transformed into a slight bias in the opposite direction. For MD-Cat, the trends for internal branches are similar to other methods, but terminal branches have a small underestimation bias that is slightly increased when using CoalBL in conditions with calibration points; as a result, it has an underestimation bias for node ages. When no calibration points are used, trends change for MD-Cat but remain similar for other methods. Similar patterns are observed when examining node ages (Fig. S8).

Increasing the number of genes tends to improve the accuracy for all methods, but it also impacts the relative accuracy of methods (Fig. 3c), with similar patterns across various metrics (Fig. S9). Setting aside the least accurate method, LSD, given only 50 genes, CoalBL tends to be *worse* than ConBL. As the number of genes increases beyond 200, CoalBL performs better regardless of the sequence length or dating method. The impact of the number of genes is perhaps the clearest with 1600bp (i.e., accurate) gene trees, where CoalBL is worse than ConBL with 50 genes but becomes better at 200, and its advantage increases going from 200 to 1000 genes.

Both tMRCA and treeness have a substantial underestimation bias with ConBL and a smaller overestimation bias with CoalBL for all sequence lengths, number of calibrations, and dating methods (Fig. 3de). For the most difficult replicates, the over/underestimation can reach 2*×* or more (Fig. S10). The trends are consistent across different number of calibrations, but using more calibrations reduces the variation in tMRCA estimations of CoalBL pipelines, while having little to no impact on ConBL-based pipelines (Fig. S10).

### Scalability experiments

We evaluate the scalability of ConBL and CoalBL dating pipelines (both run with TreePL, which was the fastest ML-based dating method in our simulations; Figs. 4, S11 and S12). On the datasets with more than 1000 taxa, ConBL fails to finish with 64GB of memory on all replicates. In conditions with up to 1000 taxa, using CoalBL is far more scalable than using ConBL. With 1000 taxa, CoalBL finishes in about a minute on average, while ConBL takes more than an hour. In this model condition, ConBL uses up to 49 GB of memory, while CoalBL uses up to 3 GB on average. In addition, CoalBL finishes in all 20 replicates of the final model condition with 10,000 species, taking 1.6 hours on average and up to 26 GB of memory. In conditions with 5,000 and 10,000 species, the dating step (TreePL) takes 76.93% and 56.15% of the total runtime and CASTLES-Pro uses less than 45 and 15 minutes on average, respectively, whereas in other conditions, more than 75% of the total runtime is used by CASTLES-Pro (Fig. S11).

**Figure 4:**
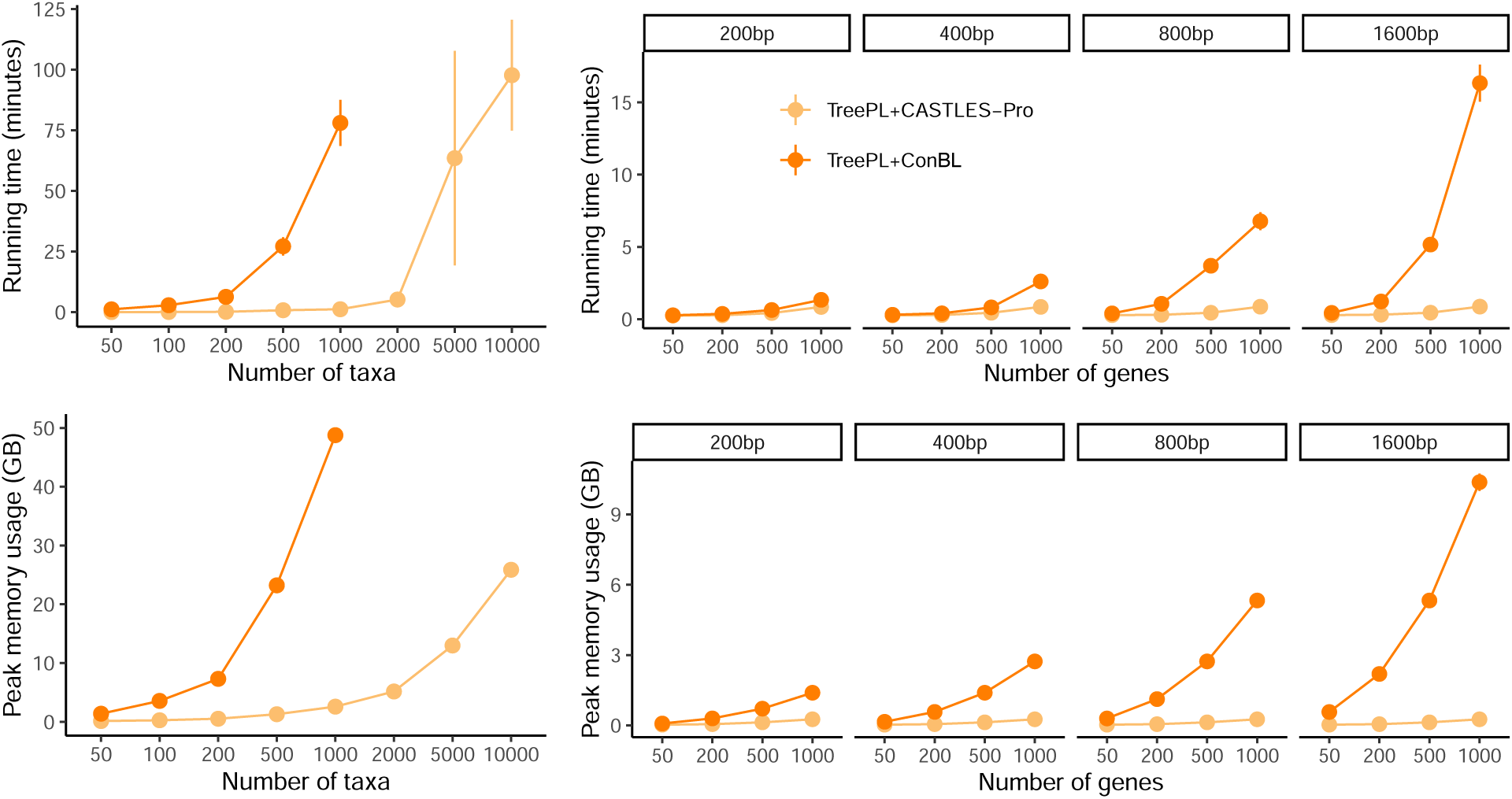
Scalability on the large (left) and S100 (right) simulated datasets, varying number of taxa, number of genes and sequence length. Results show average total runtime of branch length estimation and dating and peak memory usage for TreePL with CoalBL or ConBL in the *root-fixed* experiments. The reported runtime does not include the time spent for gene tree estimation or species tree topology estimation, as those are assumed to be available beforehand, and ConBL and CoalBL both estimate branch lengths for a fixed tree topology. ConBL fails on trees with more than 1000 taxa due to memory limit. The results are averaged over 20 replicates for the large dataset and 50 replicates for S100 dataset in each model condition. See also Figs. S11 and S12

On the S100 dataset, for all numbers of genes and all sequence lengths, CoalBL pipeline uses less time and memory than ConBL. As the number of genes or the sequence length increases, the gap between the runtime and memory usage of the two pipelines increases, so that for 1000 genes with 1600bp sequence length, CoalBL is more than an order of magnitude faster and more memory-efficient (260MB compared to 10 GB on average) than ConBL. In all cases when using ConBL, the runtime and memory usage is almost entirely dominated by the branch length estimation step using RAxML in conditions with more than 50 genes (Fig. S12b). In pipelines using CoalBL, however, TreePL takes 96%, 84%, 56% and 28% of the total runtime when using 50, 200 and 500 genes, respectively, dominating the CASTLES-Pro step.

### Suboscines: corrected species diversification rates

Using CoalBL instead of ConBL for dating the subsocines phylogeny results in substantial reductions in the length of terminal branches and ages of shallow nodes for both ASTRAL and concatenation topologies (Figs. 5, S13 and S14). Using either topology, the mean terminal branch length decreases by 5% or more for 23 out of 36 suboscines families when we use CoalBL instead of ConBL, with a marked difference in Philepittidae, Platyrinchidae and Strigopidae families with 2X decrease (Fig. S15). Similar to results in simulations, the treeness of the dated trees increases when using CoalBL instead of ConBL (from 0.4 to 0.46 for the concatenation topology and from 0.35 to 0.40 for ASTRAL).

**Figure 5:**
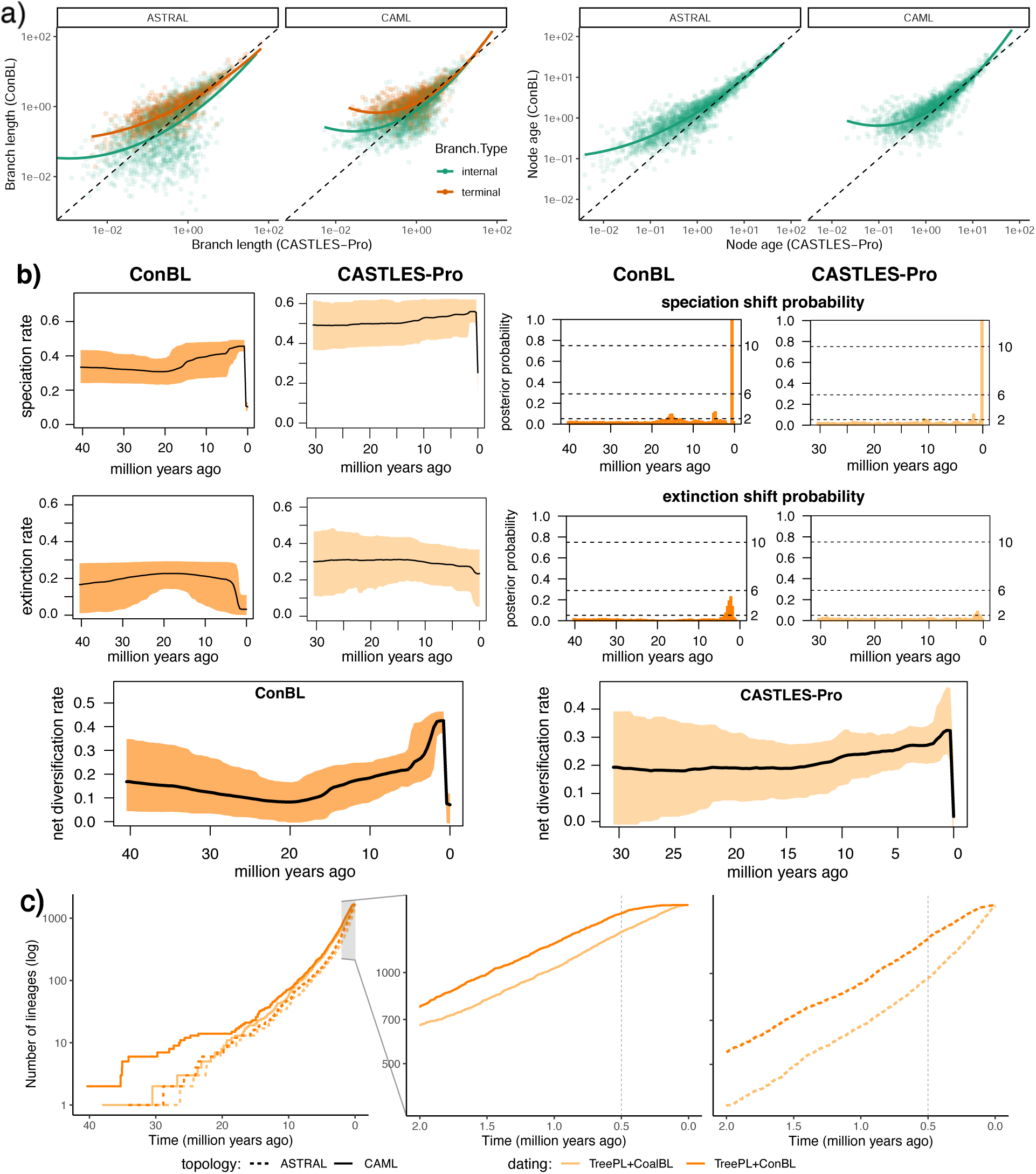
Results on the Suboscines dataset of Harvey et al. (2020). Dating is done using TreePL with four calibration points as in the original study. a) Correlation between time-unit branch lengths and node ages of the trees dated based on branch lengths of ConBL and CoalBL on the concatenation and ASTRAL topologies. b) Diversification rates, including speciation and extinction rates and speciation and extinction shift times for the concatenation topology from the original study (after removing the outgroup clade) based on branch lengths estimated using ConBL or CoalBL. c) Lineage-through-time plots (log scale *y*-axis) for the dated trees with the concatenation topology. The left plot shows the complete diversification history of the group and the right plots focuses on the final two million years. See also Figs. S13-S17.

Similar to the results reported in Harvey et al. (2020), there is a significant drop in diversification rates, in particular speciation rates, in the last 2 million years when both topology and branch lengths are estimated using concatenation (Fig. 5b). Using CoalBL instead of ConBL on the concatenation topology reduces the drop in speciation rates and almost eliminates the final drop in extinction rates. As a result, the net diversification rates are far more stable with CoalBL dating compared to ConBL: across the 30–40 million years of suboscine evolution. The trends when using the ASTRAL topology are similar, with the net diversification rates becoming much more stable up until the final 1 Mya (Fig. S16). On this topology, CoalBL fully eliminates the final drop in both speciation and extinction rates and results in almost completely constant rates across the diversification history of suboscines.

Therefore, for all three topologies, the use of CoalBL for branch length estimation reduces the severe drop in diversification rates close to the present time. These drops were likely an artifact of the overestimation bias of concatenation for terminal branch lengths. The results for the ASTRAL topology show more stable speciation, extinction, and diversification rates when using CoalBL branch lengths compared to ConBL. However, it is worth noting that Harvey et al. (2020) preferred the concatenation topology due to discrepancies with known relationships observed in the ASTRAL topology. These discrepancies are likely due to very high levels of gene tree estimation error, given short UCEs and large numbers of taxa.

The 1683 species of the suboscines dataset are distributed across 36 families and 325 genera. Examining the age of the genera with more than 10 representative species (Fig. S17a), the age estimated by CoalBL is shallower than the age estimated by concatenation for 27 out of 39 genera. Notable reductions in clade ages occur for *Pitta* (from 15.45 to 10.10 Mya) and *Gralria* (from 14.93 to 11.08 Mya). Assessing the 19 families with at least two representative genera (Fig. S17b), the dates estimated by TreePL with the CoalBL pipeline are younger than those based on ConBL in 16 out of the 19 families, with an average decrease of 18.36%.

Lineage-through-time (LTT) plots also show notable differences between ConBL and CoalBL (Fig. 5). Regardless of the topology, LTT plots for all dated trees agree with a birth-death diversification model (i.e., showing a steep upward shift in the slope of the LTT curves in semi-log scale near the present time (Helmstetter et al., 2022)) for up until the past half a million years. However, the concatenation tree dated with ConBL shows a drastic slowdown in the LTT curve right around half a million years ago and nearly halts closer to the present; the ASTRAL tree dated using ConBL also shows a slowdown but less drastically (Fig. 5c). The LTT plots of CoalBL trees do not show this slowdown with either topology and, in fact, show accelerated growth with the ASTRAL topology. This trend aligns with the significant drop in diversification rates close to the present time observed for trees dated based on ConBL but not CoalBL. The overestimation bias for terminal branch lengths (seen in simulations for ConBL but not CoalBL) can explain the unexpected drop in rate of growth in LTT plots.

### Neoavian phylogeny

We used the three most accurate ML-based dating methods in simulations (TreePL, MD-Cat, and wLogDate) to date the 363-taxon avian phylogeny of Stiller et al. (2024) where the topology was estimated using ASTRAL. Using CoalBL instead of ConBL results in shorter terminal branches and shallower recent nodes for all three dating methods (Figs. 6 and S18). This trend agrees with the results in simulations in which ConBL produced longer terminal branches than CoalBL. For internal branches, in contrast, lengths were similar (with the exception of MD-Cat, where CoalBL results in shorter internal branches). Treeness was higher for CoalBL combined with MD-Cat dating (0.291 versus 0.277), just as in simulations, but TreePL (0.271 versus 0.269) and wLogDate (0.286 vs 0.285) did not result in significant differences in treeness.

**Figure 6:**
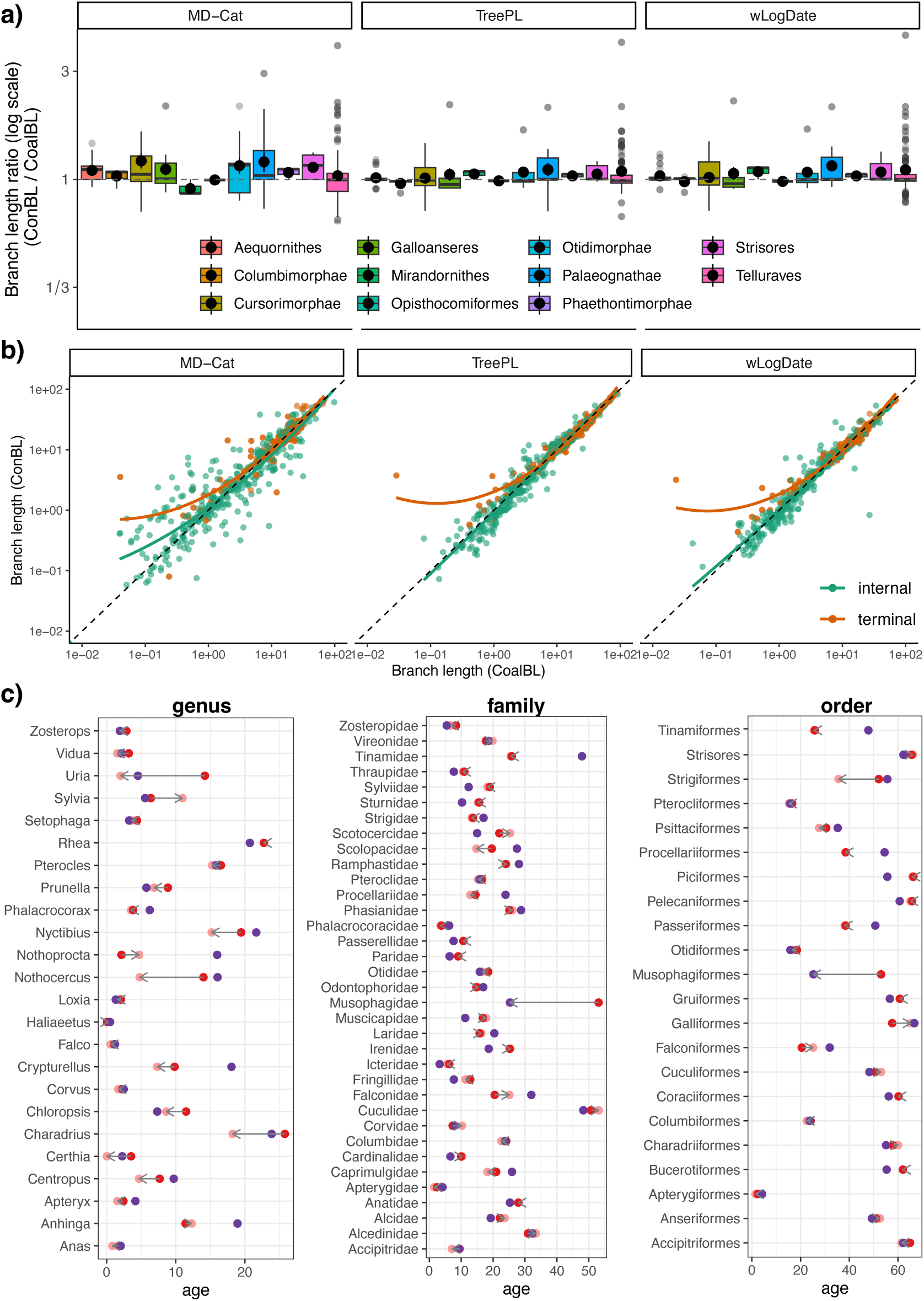
Dating the 363-taxon ASTRAL topology from the (Stiller et al., 2024) bird study using TreePL, MD-Cat and wLogDate. a) Terminal branch lengths of dated trees based on ConBL divided by terminal branch lengths of dated trees based on CoalBL in log2 scale across 11 higher order clades. b) Correlation between all branch lengths estimated by dating pipelines based on ConBL and CoalBL. c) Age of the genera that have at least two representative species and the age of families and orders with at least three representative species estimated by MD-Cat (see Figs. S21 and S22 for TreePL and wLogDate) based on CoalBL, ConBL, as well as MCMCtree analysis from the original study (which used a subset of loci used in other analyses; provided for completeness). See also Figs. S18 to S22.

Terminal branches produced by ConBL are, on average, longer than those produced by CoalBL in 9 out of 11 higher-order clades (Fig. 6) and only slightly shorter in two clades (Mirandornithes and Opisthocomiformes for MD-Cat, and Columbimorphae and Opisthocomiformes for Tree-PL, with ratios greater than 0.9). The differences are most dramatic for the hard-to-place orders Strigiformes (owls) and Apterygiformes (kiwis), as well as Falconiformes (falcons) and associated families (Fig. S20). Changes in terminal branch lengths also affect the ages of recent nodes. At least 19 out of 24 genera with two or more representative species were younger in the CoalBL results (regardless of the dating method used), and in some cases, the difference was substantial (Figs. 6, S21 and S22c). For example, for the MD-Cat tree, *Uria* goes from 14 Mya using ConBL to 2 Mya using CoalBL.

Differences in the age of families and orders, in contrast, were marginal, with a couple of notable exceptions (Fig. S20). The age of the order Musophagiformes was estimated as 25.4 Mya in the original study and 25.2 Mya in the MD-Cat+CASTLES-Pro tree, but was more than twice older in the MD-Cat+ConBL tree (53.1 Mya). The age of order Strigiformes was estimated as 35.6 Mya in the MD-Cat+CASTLES-Pro tree, which is substantially younger than its age in the MD-Cat+ConBL tree (52.1 Mya) and MCMCtree (55.6 Mya). The more stable dating of families and orders compared to genera using different dating pipelines can be explained by the observation from simulations that the overestimation bias of ConBL is mostly concentrated on terminal branches, thus affecting deeper nodes less than more recent ones (Fig. S18).

A focal point of Stiller et al. (2024) was the age of Neoaves in relation to the K–Pg extinction event (estimated at 66.04 Mya). For this deep node, the choice of the dating method matters more than whether we use ConBL or CoalBL, and the results differ slightly from the original study. Stiller et al. (2024) put the age of the MRCA of Neoaves incredibly close to the K–Pg boundary, at 67.4 Mya, and only one additional speciation event occurred prior to K–Pg boundary (though several more nodes had 95% confidence intervals that overlapped with the boundary). In all of our six dating pipelines, which unlike the original analyses do not include confidence intervals for fossils (see Material and Methods), Neoaves was dated a bit earlier (between 72.5 and 68.8 Mya, depending on the pipeline, see Table S3). Moreover, between 16 and 18 speciation events occurred right before K-Pg boundary in rapid succession in all these trees. These results, however, need to be treated with caution and underscore the importance of considering fossil dating uncertainty for dating deeper nodes of the tree.

## Discussion and future work

We introduced a scalable four-step coalescence-aware pipeline for dating species trees that uses coalescent-based methods to estimate the topology and branch lengths of the species tree from gene trees (CoalBL). Simulation analyses showed that this pipeline is substantially more scalable than pipelines using concatenation for divergence time estimation (ConBL) and easily scales to datasets with thousands of species and genes in about an hour, enabling us to date ultra-large species trees. Furthermore, dated trees obtained from this pipeline better capture species divergence times (as opposed to genic divergences) compared to pipelines using concatenation when the level of incompelte lienage sorting is at least moderate. In particular, in the presence of substantial incompelte lienage sorting, dating using concatenation has an overestimation bias for terminal branches, an issue that the coalescence-aware pipeline eliminates. Our results on two avian biological datasets corroborate results from simulations: concatenation produced longer terminal branches (especially for shorter ones), pushing genus-level nodes further back in time. The coalescence-aware pipeline reduced terminal lengths and substantially changed the age of many genera, and some families and orders (Figs. 5, 6 and S17).

An overestimation of terminal branches may compromise inferences about diversification dynamics. First, an overestimation of terminal branches may erase the signal of extinction in dated trees because extinction results in an apparent excess of recent cladogenetic events, producing an upturn in the log-scaled lineages-through-time plots (Nee et al., 1994). In contrast, empirical dated trees derived from concatenation methods not only rarely show an upturn in the LTT plot but more frequently show a downturn near the present (e.g., Cadena et al., 2020; Derryberry et al., 2011). In fact, diversification slowdowns are detected so frequently in empirical phylogenies that they have inspired a variety of hypotheses for their explanation (Moen and Morlon, 2014). Slowdowns have been attributed to incomplete species sampling (Stadler, 2009), non-random species sampling (Cusimano and Renner, 2010), DNA substitution model misspecification and thus shortening of internal branches (Revell et al., 2005), protracted speciation (Etienne and Rosindell (2012), geographic speciation (Pigot et al., 2010), ecological niche saturation (Weir, 2006), and gene tree-species tree discordance (Burbrink and Pyron, 2011). Our simulations suggest that deep coalescence of gene trees may be a major source of terminal branch overestimation, resulting in misleading inference of extinction-free slowdown in diversification dynamics. Our new methodology provides a solution to this pervasive bias and may improve studies of diversification dynamics.

Our simulation studies show that using a coalescent-aware dating is especially beneficial when few calibrations are available, where the overestimation bias of concatenation is largest. Therefore, the discordance-aware pipeline can be most useful to biologists in studies where the size of the species tree is large but few fossil calibrations (or sampling times) are available. While the four-step dating pipelines have substantially less bias than pipelines based on concatenation and reduce the overestimation bias for terminal branches, they are not completely unbiased. In particular, when gene tree estimation error or deviation from the clock is high, coalescent-based dating pipelines can have a small underestimation bias for terminal branches (Fig. 2). In addition, the four-step pipeline is more impacted by gene tree estimation error; when using very few genes or gene trees with very high gene-tree estimation error, it can be less accurate than using concatenation (Fig. 3).

For the third step of our pipeline, substitution branch length calculation from gene trees, we only considered CASTLES-Pro. This was motivated by results by Arasti et al. (2024); Tabatabaee et al. (2025) that show that it outperforms other methods in terms of branch length accuracy and scalability. However, as better methods are developed for this step, they can be used instead of CASTLES-Pro. The goal of the present paper is not to argue that CASTLES-Pro is the best method of branch length calculation, but rather, that coalescent-aware methods should be used in dating.

Although our simulation study covered a broad range of model conditions, it has limitations. In particular, we only simulated calibration points as fixed values, and used the true time of the calibrations as input to the dating methods. Future studies could improve upon this by incorporating more advanced calibration simulation protocols, such as modeling calibrations as probability densities or min-max bounds, which are used by some Bayesian and ML-based dating methods, or by simulating calibration points with associated errors. Such estimates of uncertainty may have to be adjusted to account for the effects of deep coalescence. Although this study focused solely on datasets with ILS, other sources of gene tree discordance can also affect dating and subsequent diversification analyses. For example, introgression can contribute to biases in divergence time estimates (Pang and Zhang, 2023). Furthermore, Arasti et al. (2024); Tabatabaee et al. (2025) show that using CASTLES-Pro or CASTLES-Pro +TCMM instead of concatenation can substantially improve the accuracy of SU branch lengths in the presence of sources of gene tree discordance other than ILS, such as gene duplication and loss and horizontal gene transfer. For some of these sources of discordance, specific dating methods have been developed. For example, MaxTiC (Davín et al., 2018) uses horizontal gene transfer events to estimate divergence times on a species tree. This method can be especially useful for bacterial or microbial organisms that have abundant levels of horizontal gene transfer (Koonin et al., 2001), but for which the fossil record is scarce, complicating the use of typical fossil-based dating methods. However, its application is limited to datasets with sufficient levels of horizontal gene transfer. The scalable dating pipeline introduced here can be applied more broadly. Future work should explore dating species trees on datasets that include multiple sources of gene tree discordance.

## Acknowledgments

This work is supported by the National Institute of Health (1R35GM142725) and a Natural Sciences and Engineering Research Council of Canada (NSERC) Discovery Grant (RGPIN-2018-06747). YT was supported in part by a Dissertation Completion Fellowship from the Graduate College of the University of Illinois Urbana-Champaign. This work used Expanse at San Diego Supercomputing Center through allocation ASC150046 from the Advanced Cyberinfrastructure Coordination Ecosystem Services & Support (ACCESS) program, which is supported by U.S. National Science Foundation grants #2138259, #2138286, #2138307, #2137603, and #2138296. We thank Tandy Warnow for helpful comments on earlier versions of this work. We thank Uyen Mai for helping us with running MD-Cat.

## Data availability

The datasets/scripts are available at https://github.com/ytabatabaee/coalescent-based-dating.

## Supplementary Materials

### S1 Details of the Experimental Study

#### S1.1 Methods and Software Commands

This section includes the details of the experimental study and software commands. All scripts and data used in this study are avaialable at https://github.com/ytabatabaee/coalescent-based-dating. The experiments were run on the University of Illinois campus cluster given 64GB of memory.

##### S1.1.1 Dating Methods

###### Least-square dating (LSD)

LSD (To et al., 2016) is an ML-based dating method that assumes a strict molecular clock and models the uncertainty of the substitution-unit branch lengths using a Gaussian distribution. LSD solves a convex optimization problem, where the likelihood function is defined in the form of a weighted least-squares criterion that minimizes deviation from a strict clock. LSD was run using QPD* mode described in To et al. (2016) with minimum branch length in the time-scaled tree (option -u) set to 0.001.

We used the LSD software (v2) available from https://github.com/tothuhien/lsd2 with the following command when inferring a unit-ultrametric tree without calibration points:

~~~
lsd2 -i <ROOTED - species - tree > -a 0 -z 1 -s <SEQ - length > -u 0.001 -o
<DATED - species - tree >
~~~

and the following command when using fossil calibrations specificed with -d

~~~
lsd2 -i <ROOTED - species - tree > -d <CALIB - file > -s <SEQ - length > -u 0.001 -o
<DATED - species - tree >
~~~

where the option -s specifies the sum of the lengths of the sequence alignments for all loci, and option -u

specifies the minimum branch length in the output time-scaled tree, which we set at 1e-3.

###### wLogDate

wLogDate (Mai and Mirarab, 2021) is an ML-based method for estimating divergence times that uses a similar optimization function as LSD, but it uses a log transformation before minimizing the variance of the branch rates. The advantage of wLogDate to LSD is that the log transformation results in a symmetrical penalty function, where increases or decreases in the rates have a similar effect on the optimization function, which is useful when the clock model deviates from the strict molecular clock. However, unlike LSD, wLogDate solves a non-convex optimization problem. wLogDate was run with default settings in our experiments.

We used the wLogDate software (v1.0.4) available at https://github.com/uym2/ wLogDate with the following command, where the option -t -b specifies the fossil calibrations in backward time:

~~~
python 3 launch_w Log Date . py -i <ROOTED - species - tree > -o <DATED - species - tree >
[- t <CALIB - file > -b]
~~~

###### MD-Cat

MD-Cat (Mai et al., 2024) is an ML-based dating method that uses a categorical model of rates to approximate different clock models and co-estimates rate categories and branch lengths in the unit of time using an Expectation-Maximization (EM) algorithm. In this model, the branch rates are drawn from a discrete distribution defined by *k* rate categories (default *k* = 50). The likelihood function in MD-Cat considers a parameter for each of the *k* rate categories and the tree branch lengths in time units, and the EM algorithm is used to maximize this likelihood function and estimate all branch length and rate parameters.

We used the MD-Cat software available at https://github.com/uym2/MD-Cat with the following command:

~~~
python 3 md_cat. py -i <ROOTED - species - tree > -o <DATED - species - tree > -p 10
[- t <CALIB - file > -b] [- l <SEQ - length >]
~~~

where the option -t -b specifies the fossil calibrations in backward time and option -l specifies the length of the sequence alignment used to infer the tree. In simulations, MD-Cat was run with the number of starting points set as 10 (option -p) to speed up the runtime given the large numbr of replicates, but on biological data we used it with the default 100 starting points. On the neoavian dataset we set the sequence length parameter (-l) as 40,000. Other parameters were set as their default settings.

###### TreePL

TreePL (Smith and O’Meara, 2012) is a method for estimating divergence times on large phylogenies that uses the penalized likelihood framework of Sanderson and Shaffer (2002). This optimization function allows for rate variation across different branches, with a smoothing parameter that determines the penalty for rate differences over the tree. The TreePL algorithm operates in two steps to optimize the likelihood: a greedy hill-climbing phase and a stochastic phase using simulated annealing.

We used the TreePL software (v1.0) available at https://github.com/blackrim/treePL. For the simulation study, we set the smoothing parameter to 100 and the number of sites as the sum of the lengths of the sequence alignments for all loci. For the Neornithes dataset, we set the smoothing parameter as 1000. For the suboscines biological dataset, we used the same setting as the original study. In all experiments, We used both thorough and prime flags to do a thorough analysis and determine the best optimization parameters.

###### MCMCtree

MCMCTree (Yang and Rannala, 2006) is a Bayesian method for estimating species divergence times under different molecular clock models. Unlike the ML-based methods that use fixed calibration points or min-max bounds, MCMCTree uses soft fossil calibrations with flexible probability distributions for describing the unceratainty in fossil ages. We use MCMCTree with approximate likelihood calculation (Reis and Yang, 2011; Thorne et al., 1998) that speeds up the calculation of likelihood function during the MCMC sampling for large alignments and trees. For the *root-unfixed* experiments in the simulation studies, where the age of the root is not one of the calibration points, we set the soft bound for root age for MCMCTree as 10 + *T* where *T* is the true root age.

We used the MCMCtree software from PAML (v4.10.7) package Yang (2007) available at http://abacus.gene.ucl.ac.uk/software/paml.html under the GTR+Γ model. Most of the parameters in our analysis were taken from the MCMCtree anlaysis in Stiller et al. (2024). We set the calibration bounds as (x-0.01,x+0.01) where x is the exact calibration point. We passed the concatenation of all gene sequences as a single partition to MCMCtree (specified with ndata). For nodes without calibrations, we used a birth-death process with *λ* = *γ* = 1*, ρ* = 0.1 to get an approximately uniform kernel. We used a relaxed clock model where rates are log-normally distributed across branches and a gamma-Dirichlet prior on rates. For the simulation study, during the MCMC sampling, samples were taken every 100 steps with a total number of iterations of 5,050,000 where the first 50,000 steps were treated as burn-in. The following shows all the parameters used for running MCMCtree.

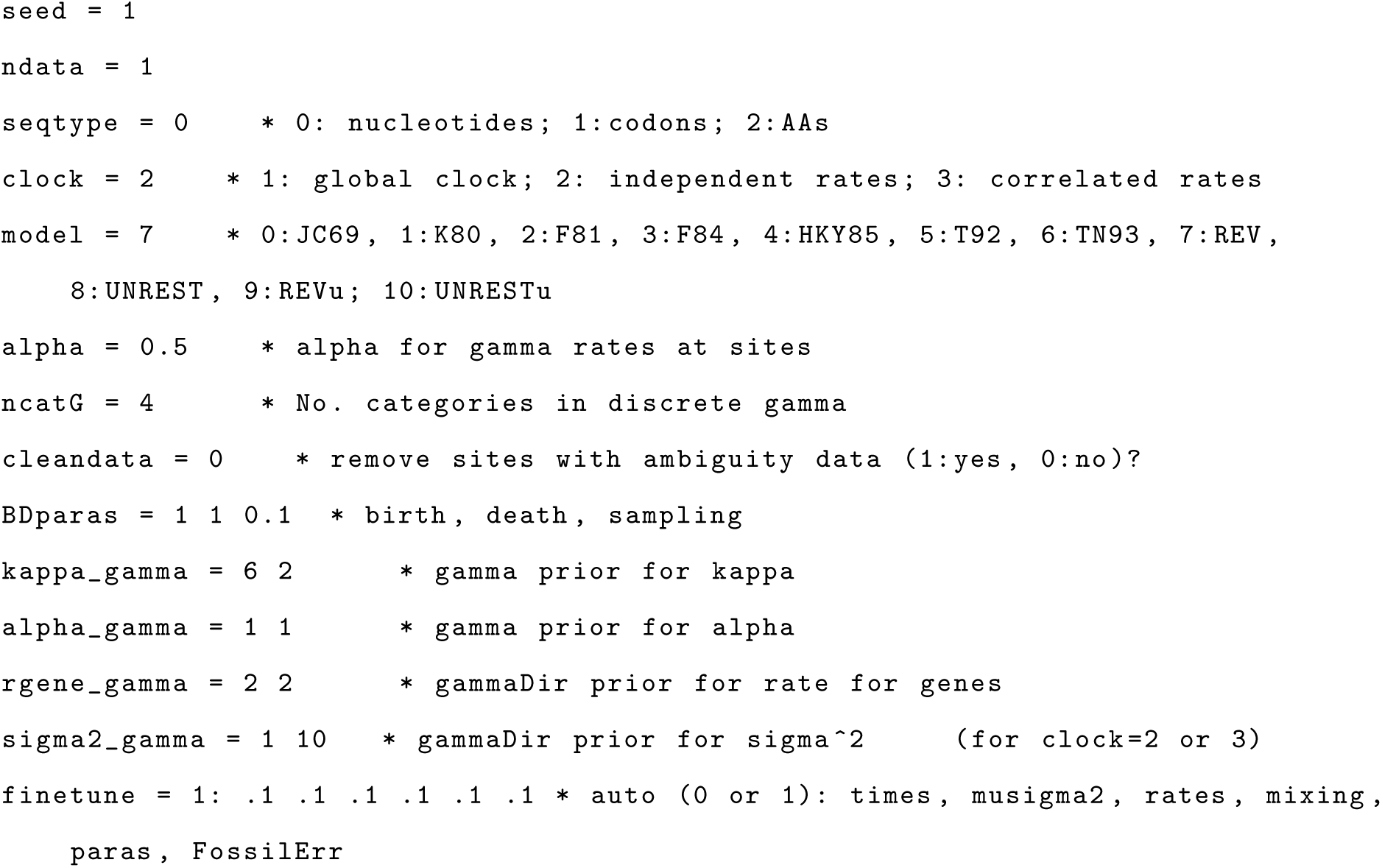
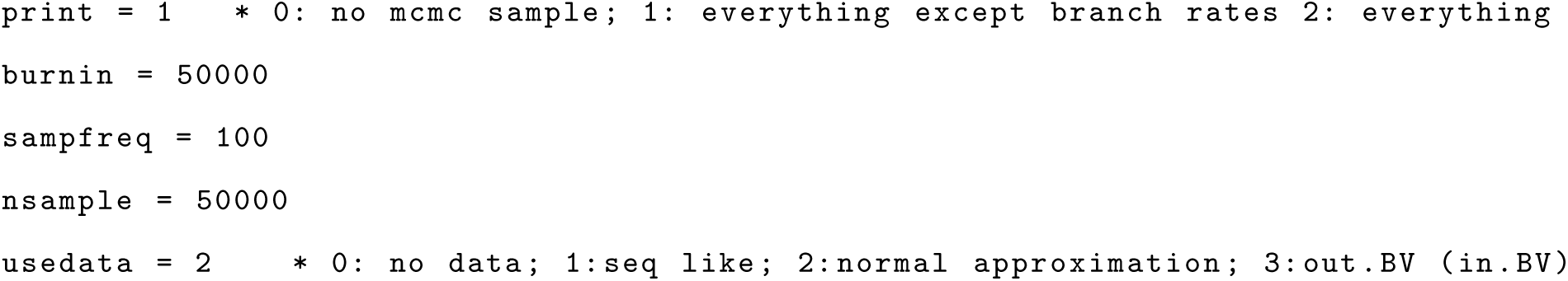

#### S1.1.2 Branch length estimation

- **CASTLES-Pro** We used CASTLES-Pro as implemented in the species tree estimation software ASTER (v1.19.3.5) available at https://github.com/chaoszhang/ASTER to estimate SU branch lengths. To estimate lengths on a fixed species tree topology (specified with the option -C -c), we used the following command:

~~~
astral4 -i <GENE - tree - path > -C -c <SPECIES - tree - topology > -o <OUTPUT - path > -- root <OUTGROUP - name > -- genelength <GENE - sequence - length >
~~~

where --root specifies the outgroup name (if known) and --genelength specifies the average gene sequence length (default: 1000bp).

- **Concatenation.** We used RAxML (v8.2.12) that is available at https://github.com/stamatak/standard-RAxML to estimate substitution-unit branch lengths on a fixed species tree topology (with the option

-f -e) using a concatenated sequence alignment. We used the following command:

~~~
raxmlHPC - PTHREADS -f e -t < species_tree_path > -m GTRGAMMA -s < alignment_path >
-n RES -p 4321 -T 16
~~~

**Table S1:** Parameters used in SimPhy simulation of large dataset.

| Arg. | Description | Value |
| --- | --- | --- |
| RS | Number of replicates | 20 |
| RL | Number of loci | 1000 |
| RG | Number of genes | 1 |
| ST | Maximum tree length | LogNormal(16,1) |
| SL | Number of taxa | 50-10,000 |
| SB | Speciation rate | LogNormal(1.0e-7,1.0e-6) |
| SD | Extinction rate | LogNormal(1.0e-7,SB) |
| SP | Global population size | Uniform(20000,2000000) |
| SU | Global substitution rate | LogNormal(-17.27461, 0.6931472) |
| HS | Species-specific branch rate heterogeneity modifiers | LogNormal (1.5,1) |
| HL | Gene-family-specific rate heterogeneity modifiers | LogNormal (1.551533,0.6931472) |
| HG | Gene by lineage specific rate heterogeneity modifiers | LogNormal (1.5,1) |
| HH | Gene-by-lineage-specific locus tree parameter | 1 |
| SO | Outgroup branch length relative to half the tree length | 1 (with outgroup) |
| CS | Random number generator seed | 14907 |

**Table S2:** Empirical statistics of the large simulated dataset. AD refers to average RF distance between the model species tree and true gene trees, and GTEE refers to average RF distance between true and estimated gene trees. The number of replicates in each model condition is 20 and the number of genes is 1000.

| Number of taxa | Number of calibrations | AD | GTEE |
| --- | --- | --- | --- |
| 50 | 3 | 35.98% | 28.84% |
| 100 | 5 | 31.22% | 29.01% |
| 200 | 7 | 45.11% | 39.08% |
| 500 | 11 | 39.33% | 34.51% |
| 1000 | 15 | 39.73% | 36.89% |
| 2000 | 22 | 51.06% | 42.21% |
| 5000 | 35 | 35.53% | 36.44% |
| 10000 | 50 | 50.18% | 43.83% |

### S1.2 Simulated Datasets

We used a modified version of the simulation software SimPhy that generates species trees with SU branch lengths to generate the large ILS dataset. We used the following command, where the parameters are taken from Table S1.

./ simphy - rs " $s" - rl f:" $g" - rg 1 - sb lu :0 .0000001,0 .000001 - sd lu :0.0000001, sb

- st ln :16,1 - sl f:" $sp " - so f:1 - si f:1 - sp u: $min_pp, $max_pp - su ln : -17.27461,

0 .6931472 - hh f:1 - hs ln :1.5,1 - hl ln :1 .551533,0 .6931472 - hg ln :1.5,1 - cs 14907

-v 3 -o $sp - ot 0 - op 1 - od 1 > log_$sp . txt

## S2 Additional Results

**Figure S1:**
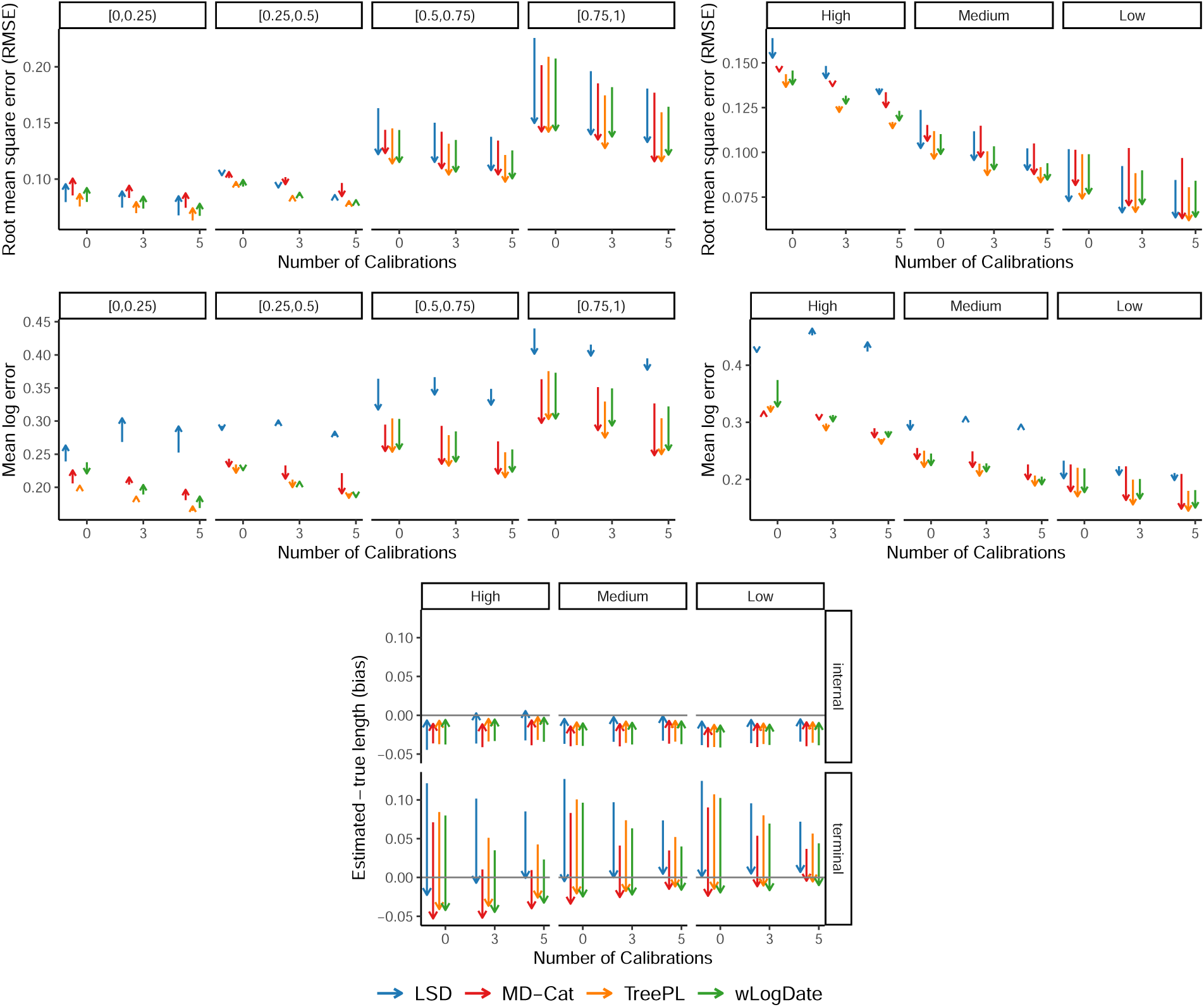
(MVroot, Branch length) Reduction (or increase) in height-normalized RMSE, mean log error and bias of branch lengths in time units using CASTLES-Pro instead of ConBL for branch length estimation in different ML-based dating pipelines. For each dating method, the direction of the arrow is from average error in the final dated tree when branch length estimation is done using ConBL to the average error when using CASTLES-Pro for branch length estimation. The results are shown on the 30-taxon simulated datasets for conditions without outgroup and in the *root-fixed* experiments.The number of genes is 500 and the number of replicates is 100. In the left panels of the top two rows, the columns show different levels of ILS and in the right panels they show different levels of deviation from the clock.

**Figure S2:**
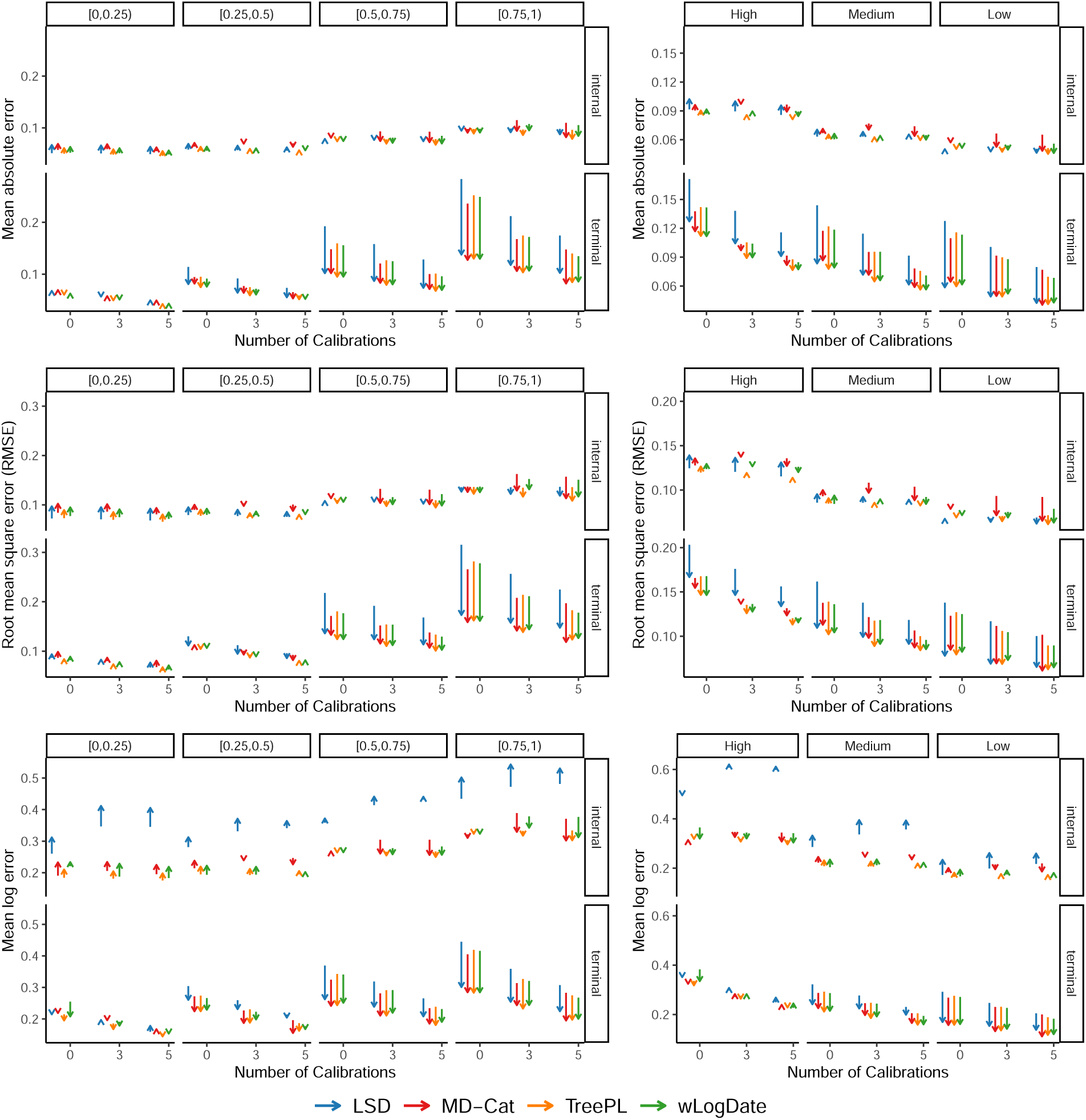
(MVroot, Branch length) Reduction (or increase) in height-normalized mean absolute error, RMSE and mean log error of terminal or internal branch lengths in time units using CASTLES-Pro instead of ConBL for branch length estimation in different ML-based dating pipelines. For each dating method, the direction of the arrow is from average error in the final dated tree when branch length estimation is done using ConBL to the average error when using CASTLES-Pro for branch length estimation. The results are shown on the 30-taxon simulated datasets for conditions without outgroup and in the *root-fixed* experiments.The number of genes is 500 and the number of replicates is 100. In the left panels, the columns show different levels of ILS and in the right panels they show different levels of deviation from the clock.

**Figure S3:**
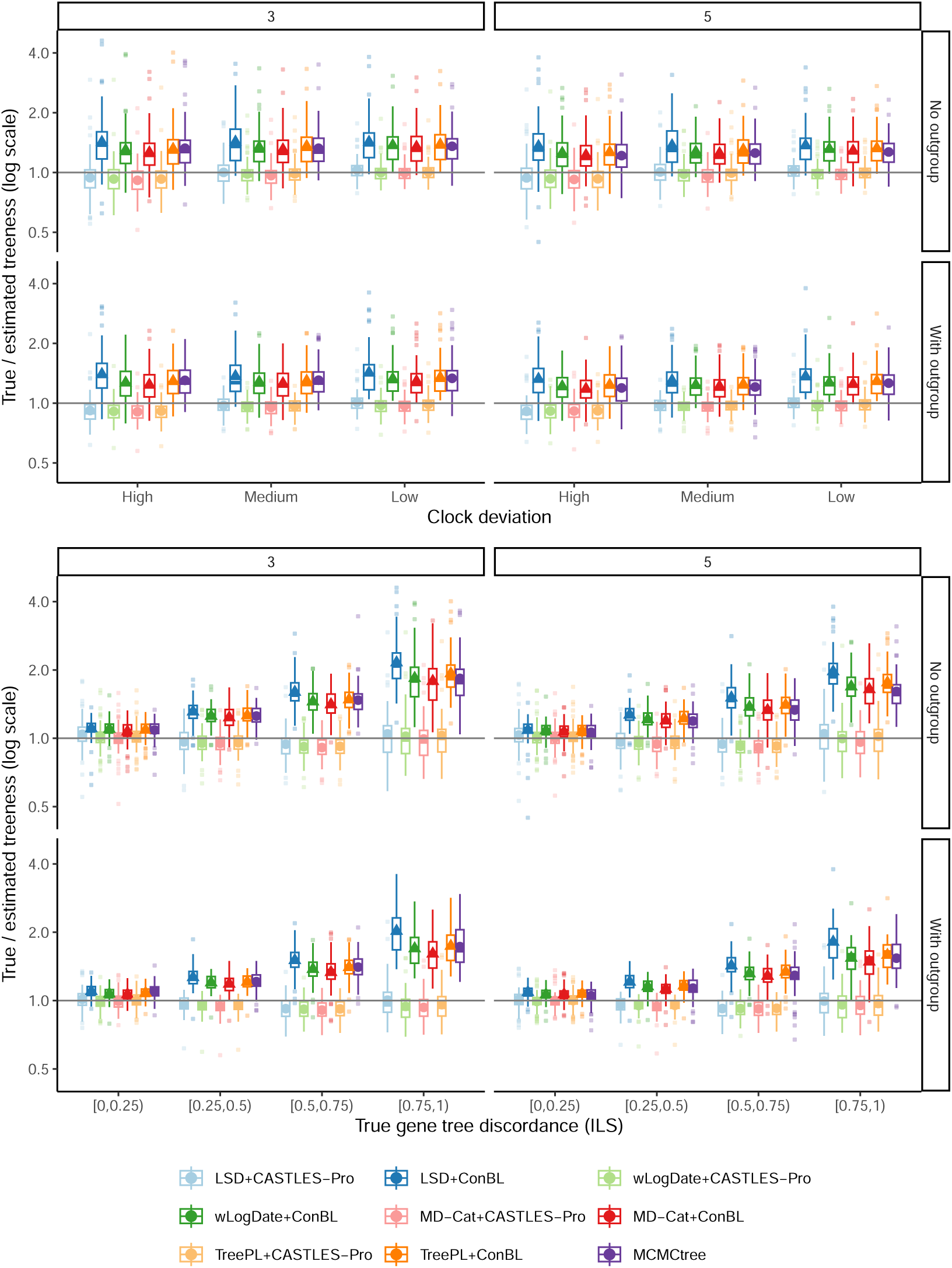
(MVroot, Treeness) True treeness (sum of internal branch lengths divided by sum of all branch lengths) divided by estimated treeness in log2 scale for trees estimated using different dating pipelines on the 30- taxon simulated datasets in the *root-fixed* experiments. The columns show the number of calibrations and the rows show conditions with or without outgroup. The number of genes is 500 and the results show mean and standard deviation in addition to boxplots across 100 replicates.

**Figure S4:**
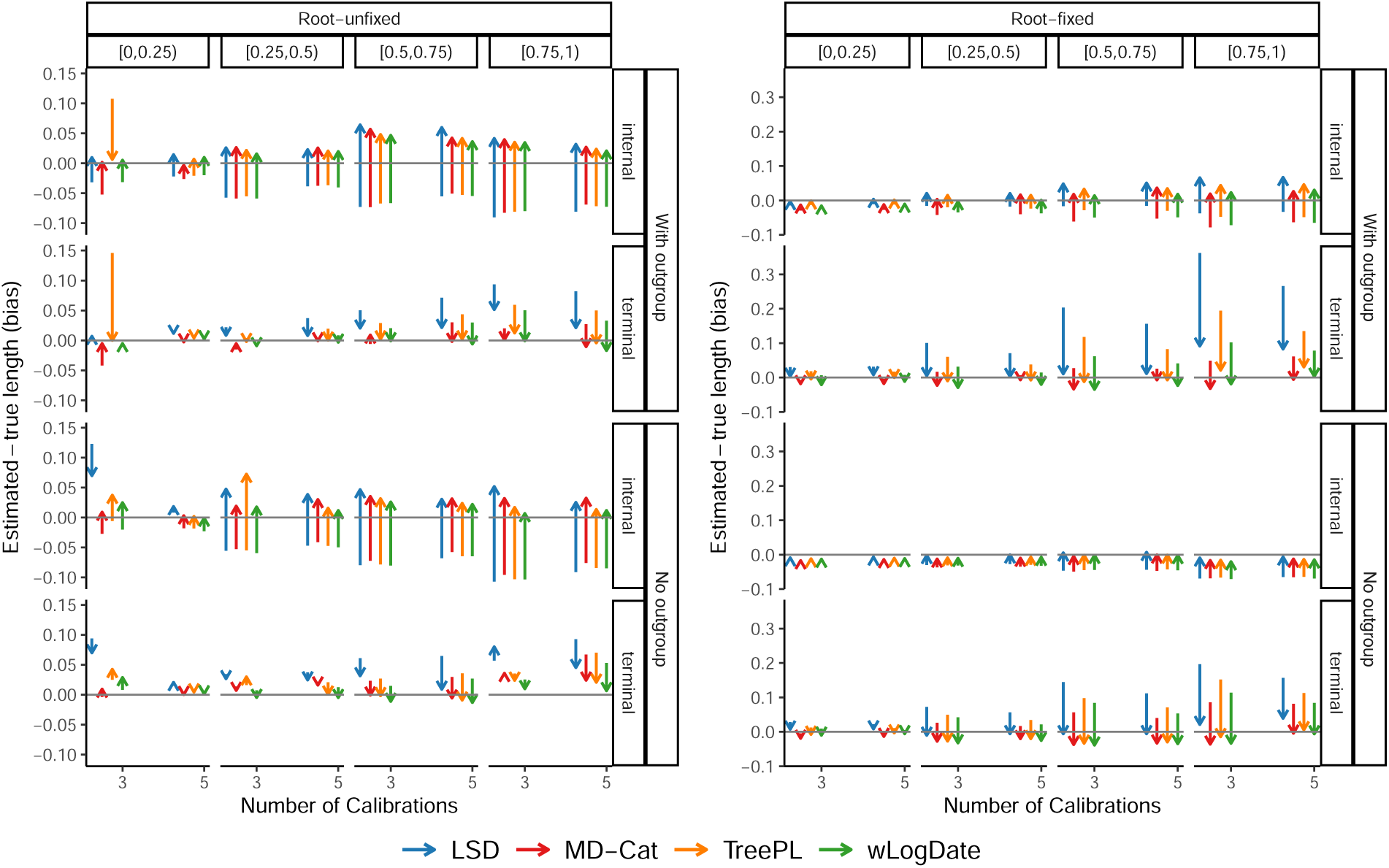
(MVroot, root-fixed vs root-unfixed) Reduction (or increase) in bias of branch lengths in time units using CASTLES-Pro instead of ConBL for branch length estimation in different ML-based dating pipelines. In conditions with outgroup, the error is calculated after excluding the outgroup. For each dating method, the direction of the arrow is from average bias in the final dated tree when branch length estimation is done using ConBL to the average bias when using CASTLES-Pro for branch length estimation. The results are shown on the 30-taxon simulated datasets for conditions with or without outgroup and in the *root-fixed* or *root-unfixed* experiments. The number of genes is 500 and the number of replicates is 100. The panels show different levels of ILS.

**Figure S5:**
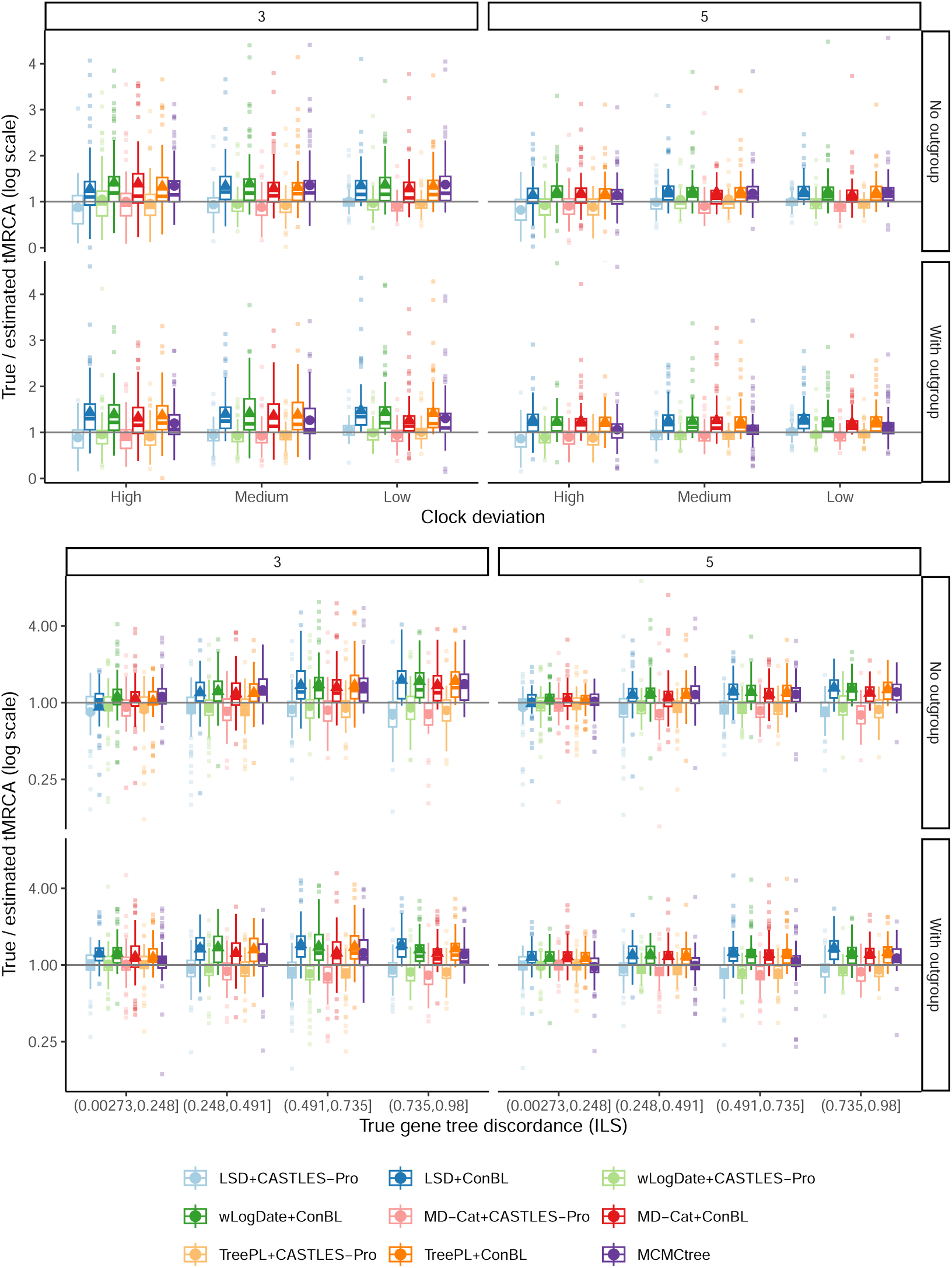
(MVroot, tMRCA) True tMRCA (time of the most recent common ancestor) of the ingroup divided by estimated tMRCA in log2 scale for trees estimated using different dating pipelines on the 30-taxon simulated dataset in the *root-unfixed* experiments. The columns show the number of calibrations and the rows show conditions with or without outgroup. The number of genes is 500 and the results show mean and standard deviation in addition to boxplots across 100 replicates.

**Figure S6:**
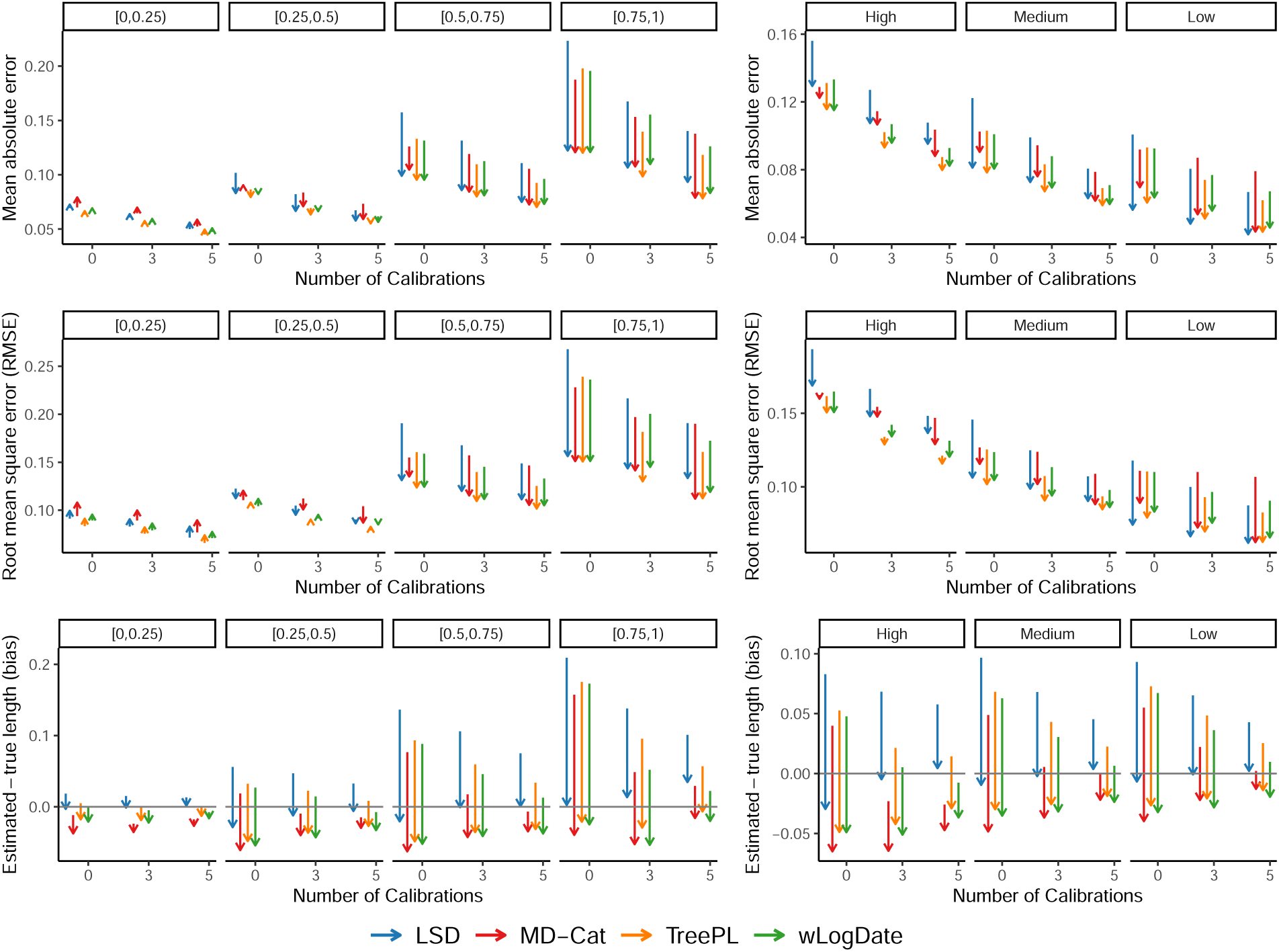
(MVroot, Node age) Reduction (or increase) in height-normalized mean absolute error, RMSE and bias of node ages using CASTLES-Pro instead of ConBL for branch length estimation in different ML-based dating pipelines. The direction of time is considered backward, so that all extant (terminal) taxa have age 0 and internal nodes have non-zero ages. For each dating method, the direction of the arrow is from average error in the final dated tree when branch length estimation is done using ConBL to the average error when using CASTLES-Pro for branch length estimation. The results are shown on the 30-taxon simulated datasets for conditions without outgroup and in the *root-fixed* experiments.The number of genes is 500 and the number of replicates is 100. In the left panels, the columns show different levels of ILS and in the right panels they show different levels of deviation from the clock.

**Figure S7:**
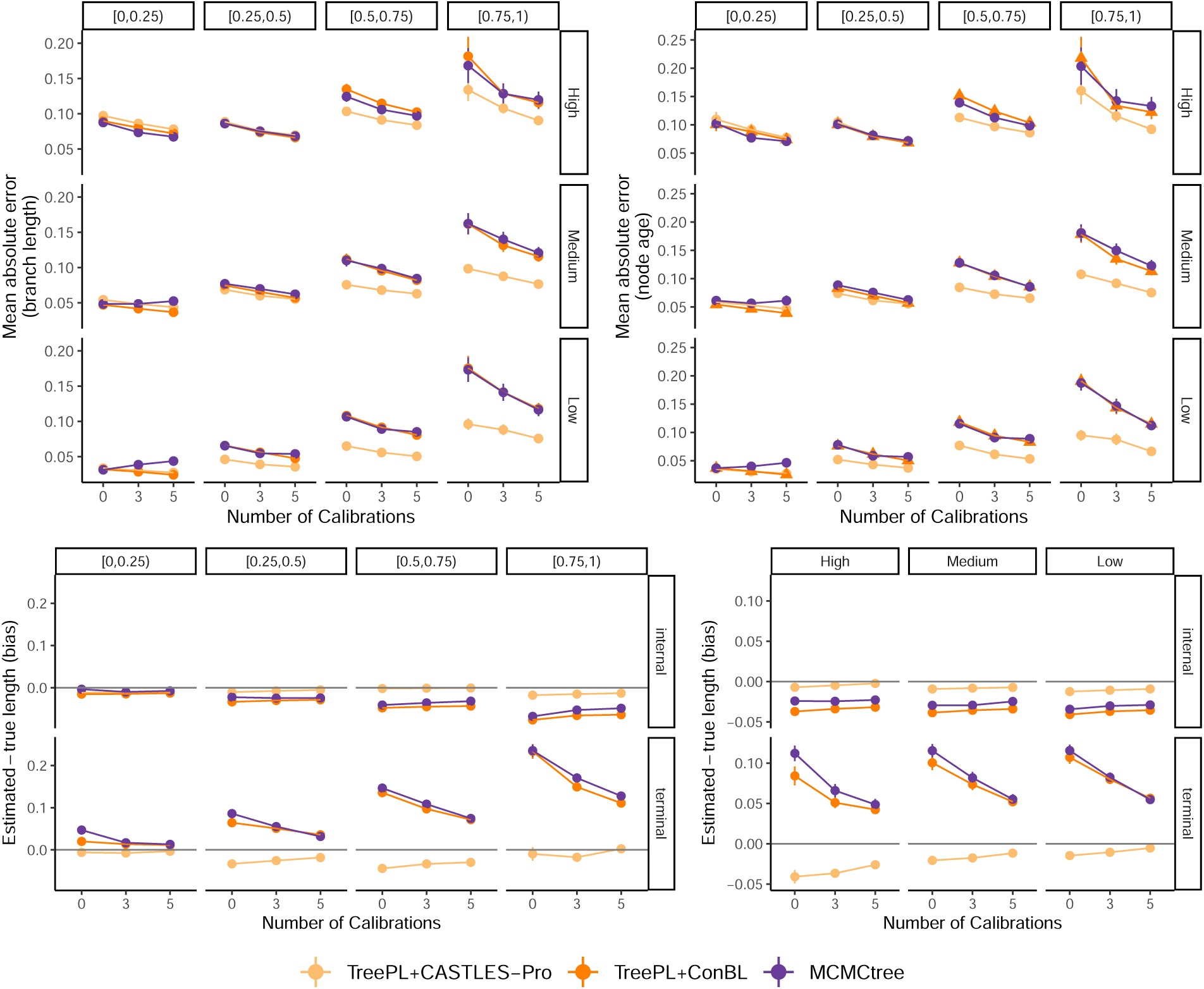
(MVroot, MCMCtree) Comparison between MCMCtree, TreePL+CASTLES-Pro and TreePL+ConBL in terms of height-normalized mean absolute error and bias of branch lengths and node ages in time unit on the 30-taxon simulated datasets for conditions without outgroup and in the *root-fixed* experiments. The number of genes is 500 and the results show mean and standard deviation across 100 replicates. For the bias plots, the left panel shows different levels of ILS and the right panel show different levels of deviation from the clock.

**Figure S8:**
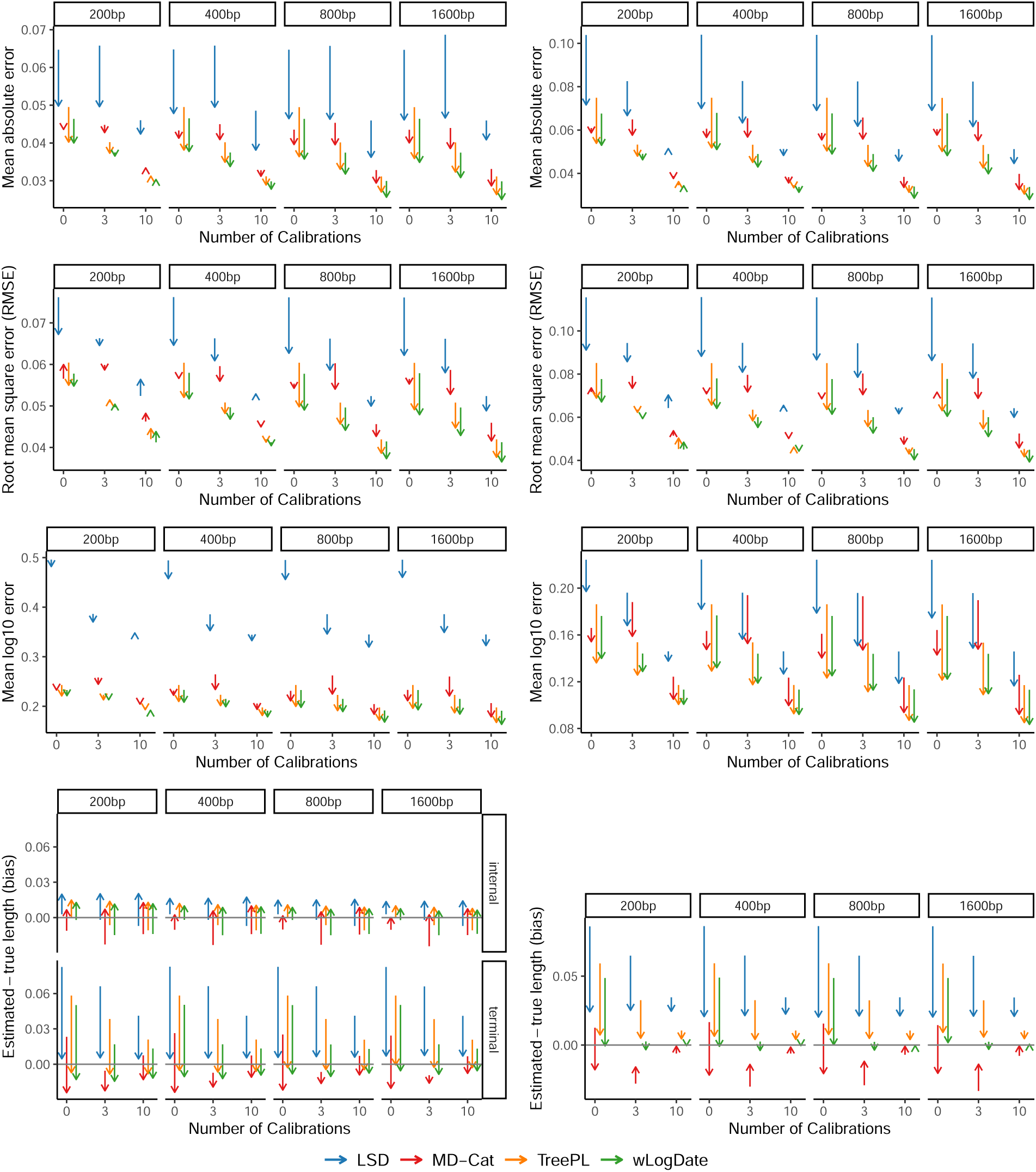
(S100, Branch length and node age) Reduction (or increase) in height-normalized mean absolute error, RMSE, mean log error and bias of branch lengths (left) and node ages (right) in time units using CASTLES-Pro instead of ConBL for branch length estimation in different ML-based dating pipelines on the 100-taxon simulated datasets in the *root-fixed* experiments. For node ages, the direction of time is considered backward, so that all extant (terminal) taxa have age 0 and internal nodes have non-zero ages. For each dating method, the direction of the arrow is from average error in the final dated tree when branch length estimation is done using ConBL to the average error when using CASTLES-Pro for branch length estimation. The number of genes is 1000 and the average level of ILS is 46% AD.

**Figure S9:**
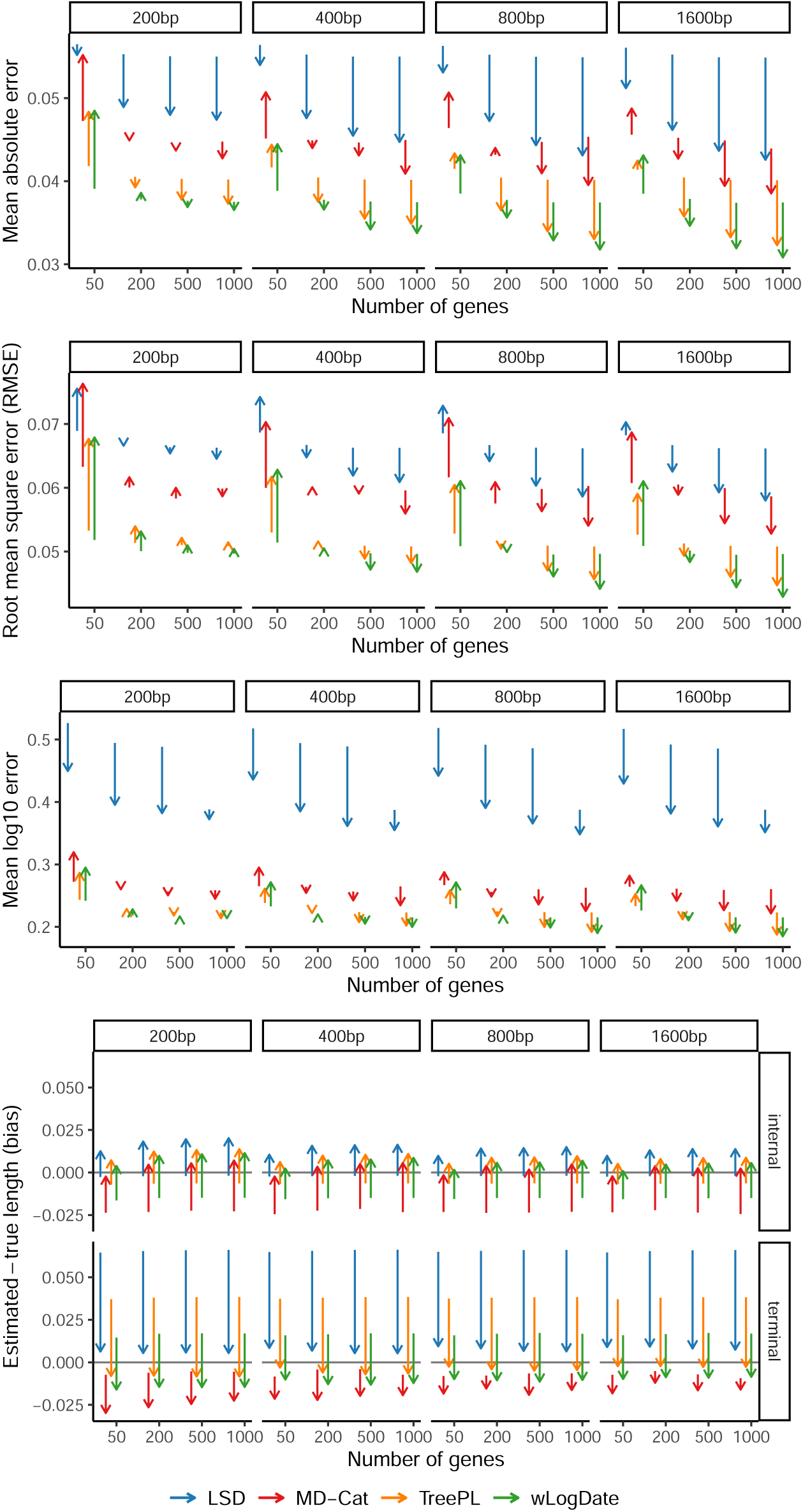
(S100, Branch length) Reduction in height-normalized mean absolute error, RMSE, mean log error and bias of branch lengths in time units estimated using different ML-based dating pipelines on the 100-taxon simulated datasets in the *root-fixed* experiments. The number of genes vary between 50 to 1000 (default: 1000) and the average level of ILS is 46% AD. For each dating method, the direction of the arrow is from average error in the final dated tree when branch length estimation is done using ConBL to the average error when using CASTLES-Pro for branch length estimation. The number of calibrations is 3.

**Figure S10:**
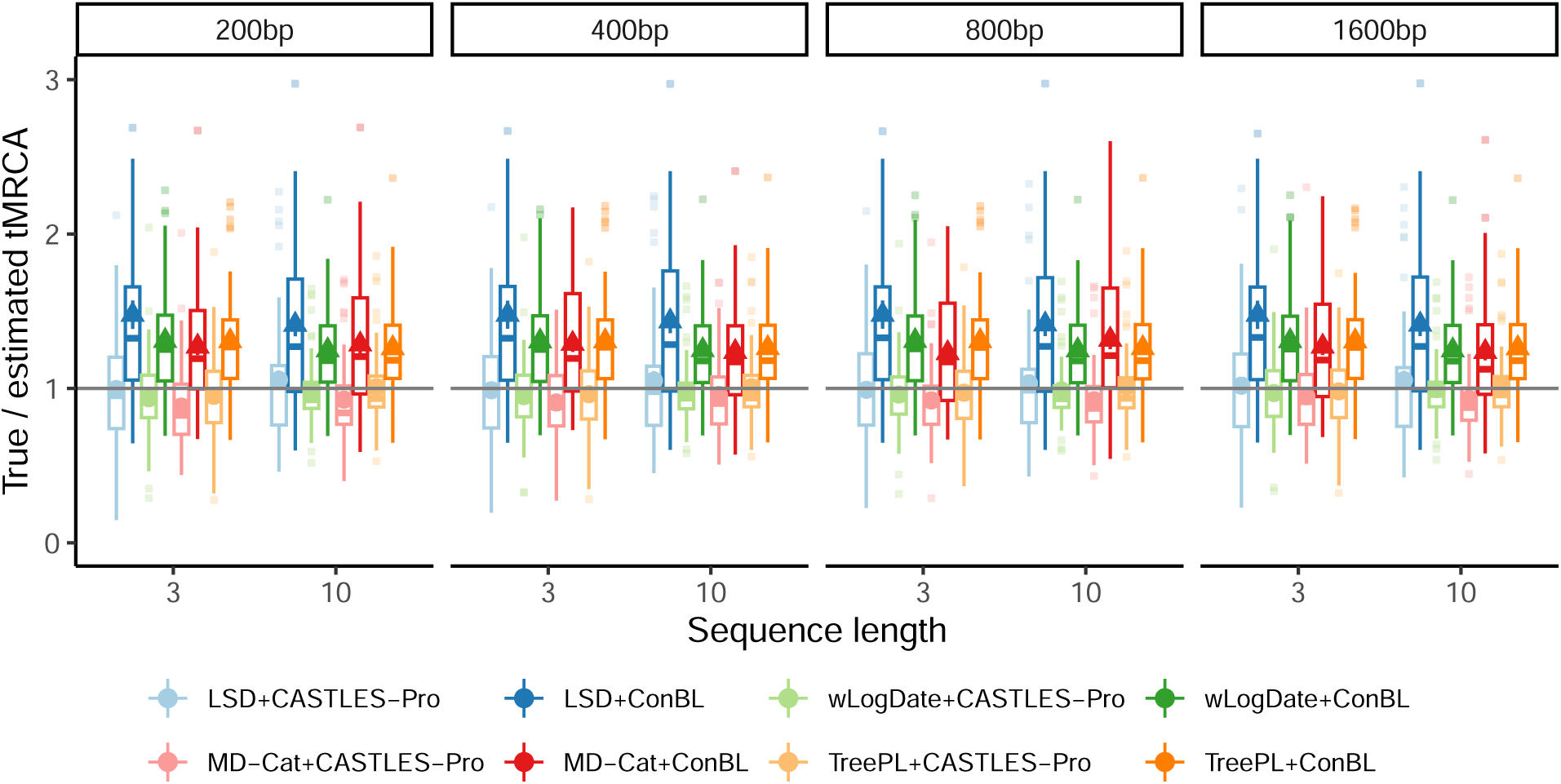
(S100, tMRCA) True tMRCA (time of the most recent common ancestor) divided by estimated tMRCA for trees estimated using different dating pipelines on the 100-taxon simulated datasets in the *root-unfixed* experiments. The columns show the sequence length. The number of genes is 1000 and the results show mean and standard deviation in addition to boxplots across 50 replicates.

**Figure S11:**
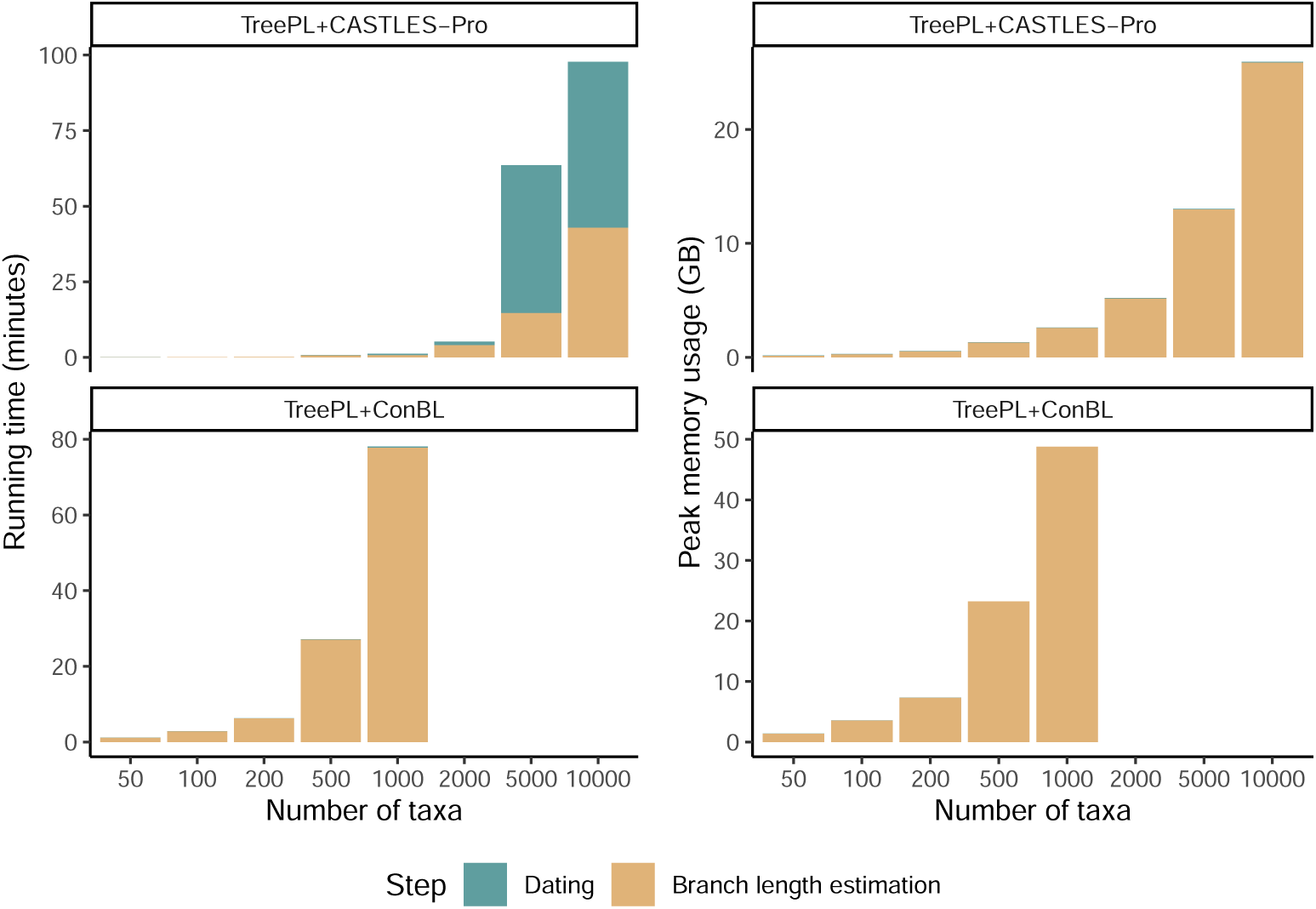
(Large dataset, time and memory) Runtime and memory seperately reported for the branch length estimation and dating steps on the large dataset. The reported runtime does not include the time spent for gene tree estimation or species tree topology estimation, as these are assumed to be available beforehand. The results are averaged over 20 replicates in each model condition. ConBL fails on trees with more than 1000 taxa due to memory limit.

**Figure S12:**
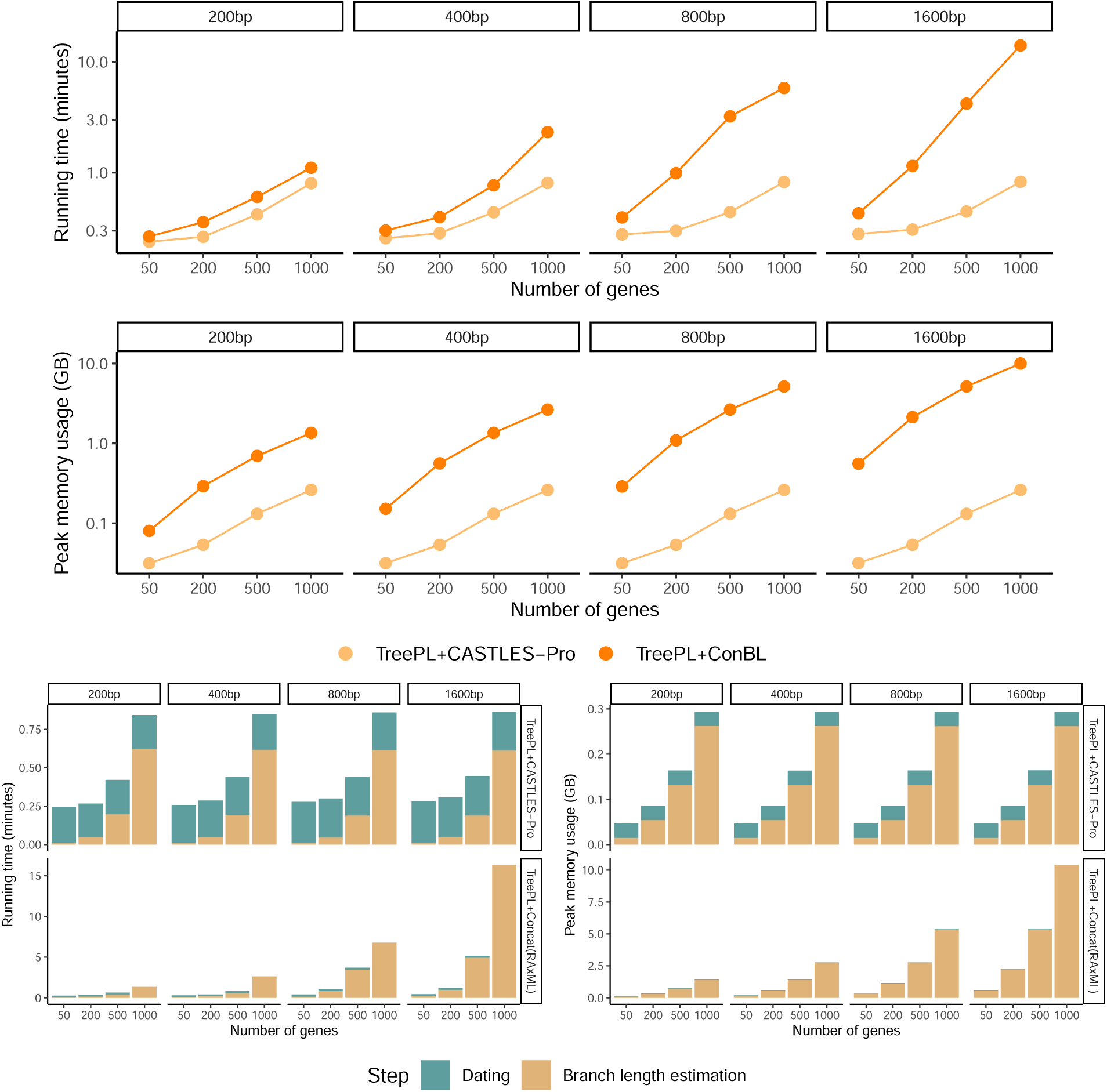
(S100, time and memory) (top) Average total running time of branch length estimation and dating in minutes and peak memory usage of the dating pipeline in gigabytes for TreePL with CASTLES-Pro or ConBL on the 100-taxon simulated datasets in the *root-fixed* experiments. The y-axes are shown in log-scale. (bottom) Runtime and memory seperately reported for the branch length estimation and dating steps. The number of calibrations is 3. The reported runtime does not include the time spent for gene tree estimation or species tree topology estimation, as these are assumed to be available beforehand. The results are averaged over 50 replicates in each model condition.

**Figure S13:**
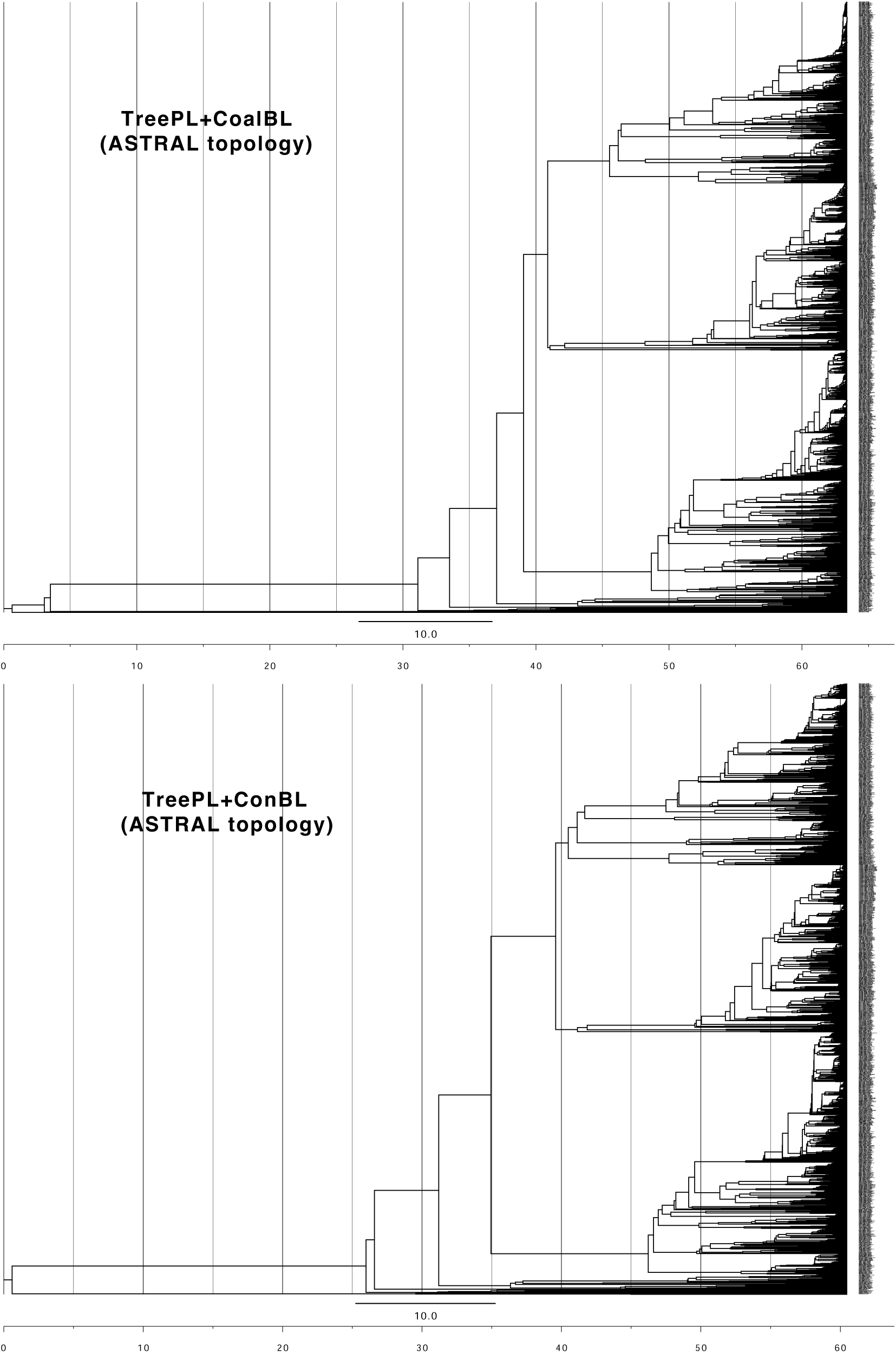
Suboscines time trees dated with TreePL using CASTLES-Pro or ConBL SU branch lengths drawn on the ASTRAL topology from the original study in Harvey et al. (2020). The number of species is 1683, and the number of gene trees is 2389. Gene trees are resolved and are estimated from the minimally filtered T400F alignments. TreePL is run using four calibration points, as in the main study.

**Figure S14:**
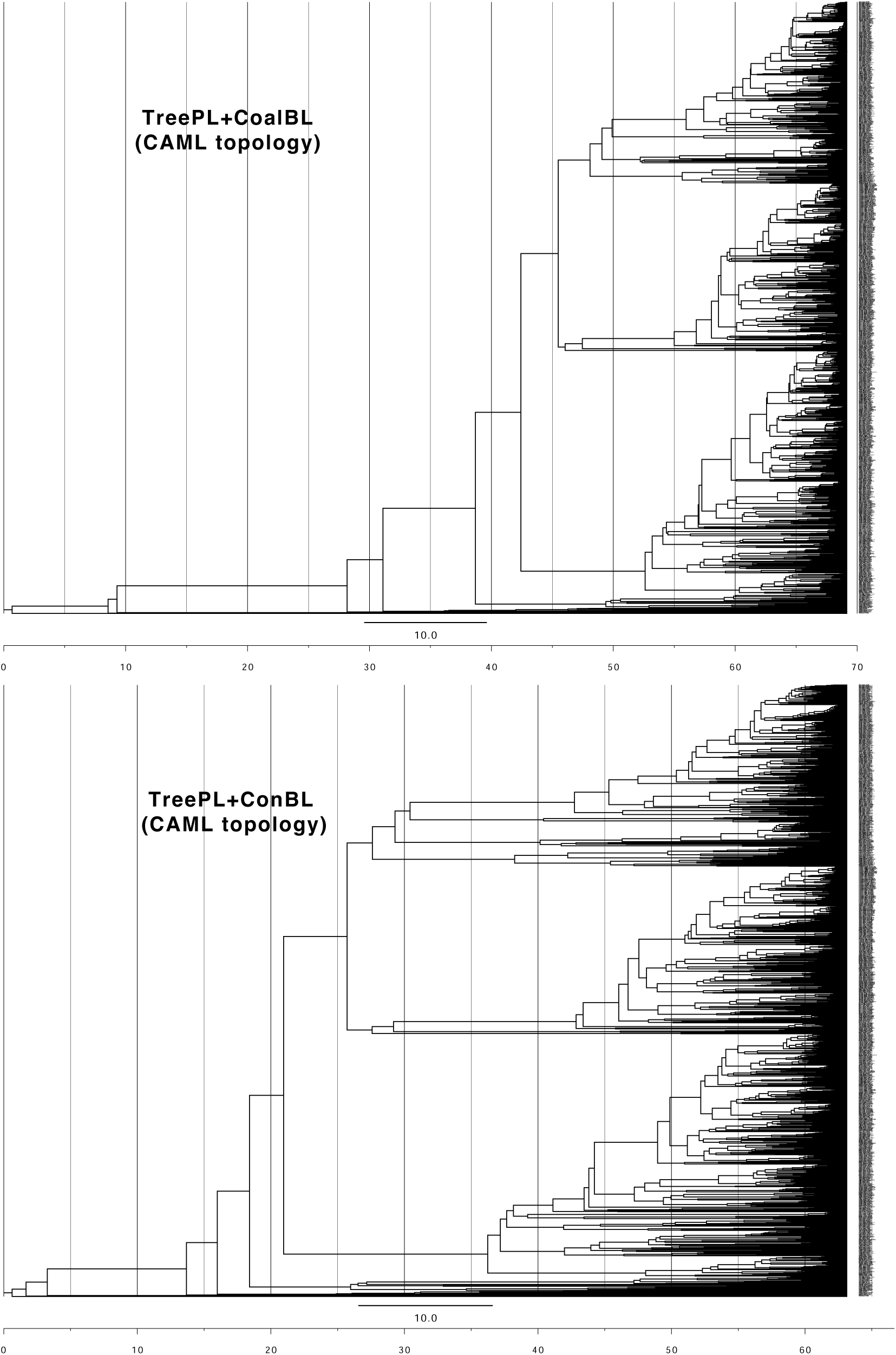
Suboscines time trees dated with TreePL using CASTLES-Pro or ConBL SU branch lengths, drawn on the concatenation topology from the original study in Harvey et al. (2020). The number of species is 1684, and the number of gene trees is 2389. Gene trees are resolved and are estimated from the minimally filtered T400F alignments. TreePL is run using four calibration points, as in the main study.

**Figure S15:**
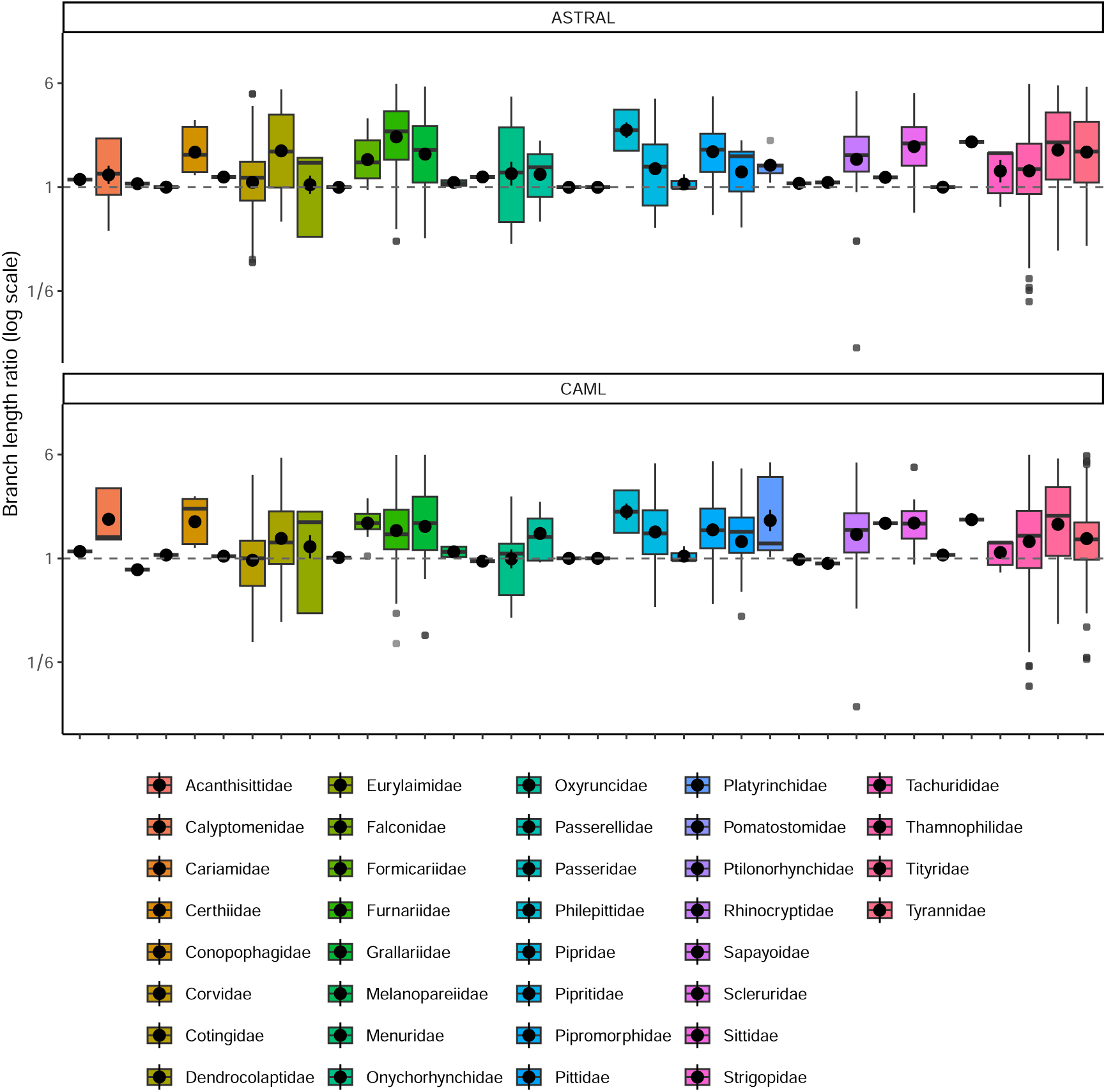
Dating the suboscines phylogeny (Harvey et al., 2020) using TreePL. The panels show the two tree topologies from the main study, estimated using concatenation and ASTRAL. The y-axis shows terminal branch lengths of TreePL+ConBL divided by terminal branch lengths of TreePL+CASTLES-Pro in log2 scale, and the x-axis shows different suboscine families. The number of species is 1683 and the number of gene trees is 2389. Gene trees are resolved and are estimated from the minimally filtered T400F alignments. TreePL is run using four calibration points as in the original study. Dotted line indicates 1.

**Figure S16:**
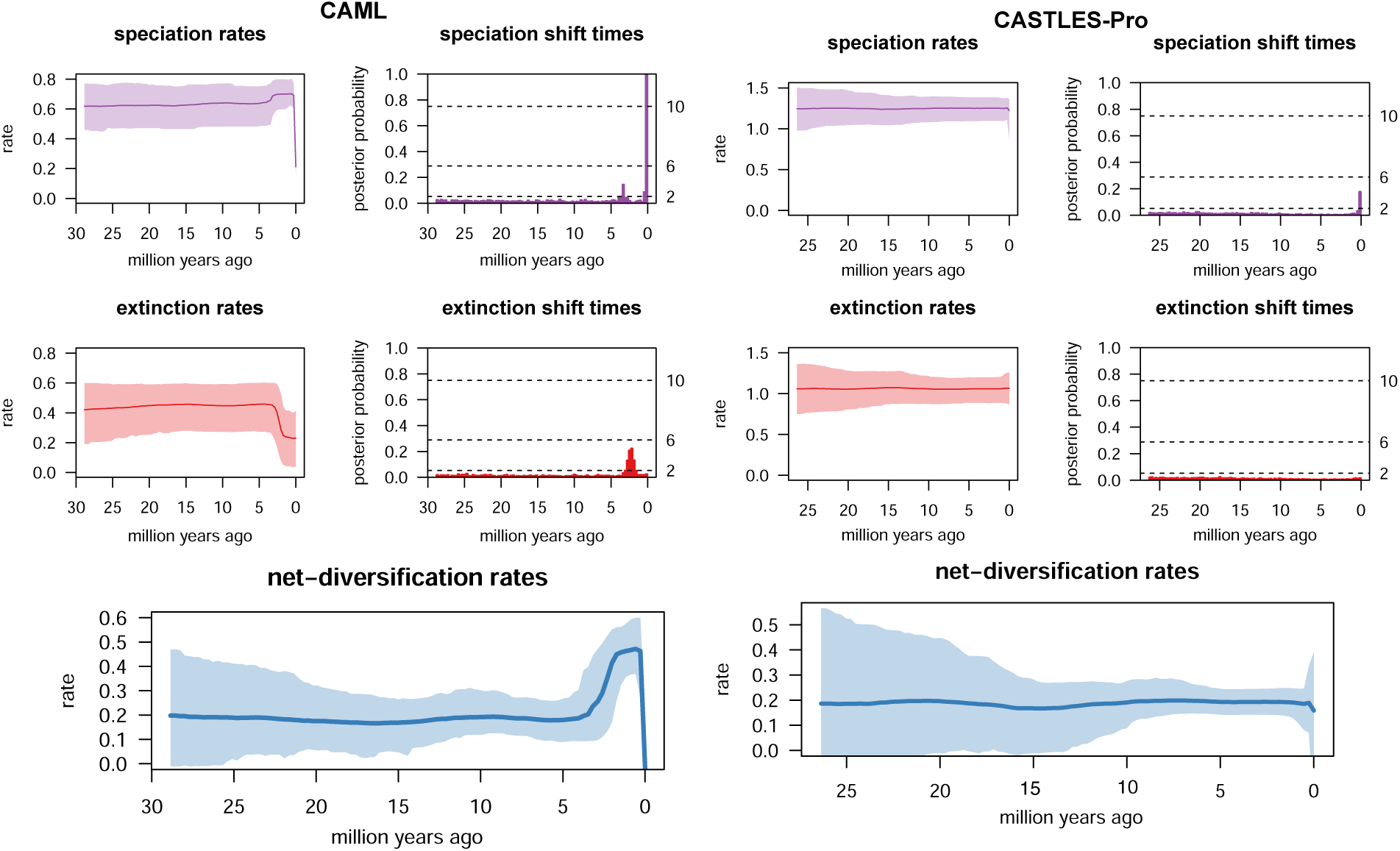
Diversification analysis on the suboscines dataset of Harvey et al. (2020). Dating is done using TreePL with four calibration points, as in the original study. The plots show diversification rates, including speciation and extinction rates and speciation and extinction shift times for the ASTRAL topology from the original study (after removing the outgroup clade) based on branch lengths estimated using ConBL (labeled CAML) or CoalBL (labeled CASTLES-Pro).

**Figure S17:**
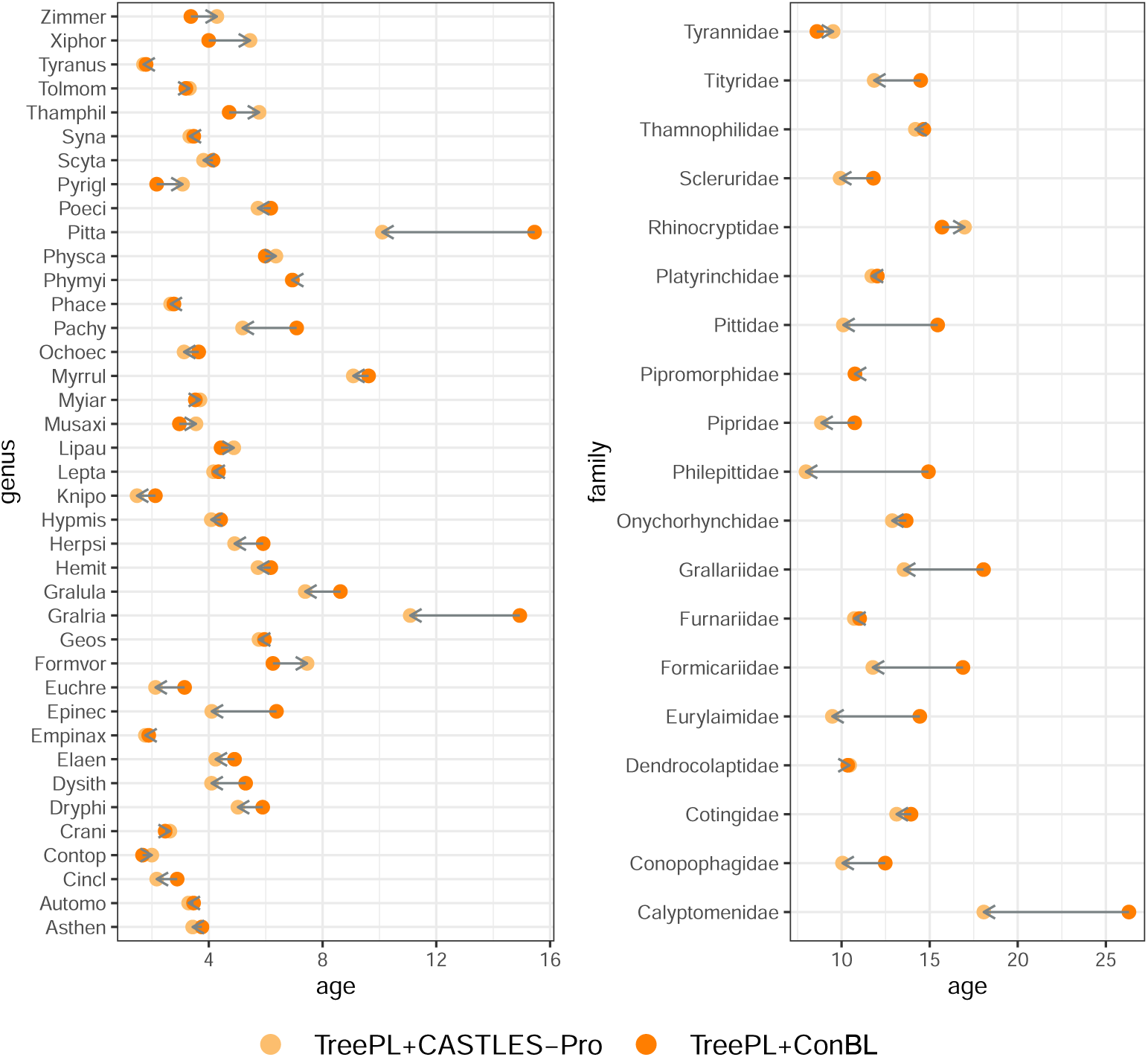
Dating the suboscines phylogeny (Harvey et al., 2020) using TreePL on the concatenation topology from the original study. Age of the genera that have more than 10 representative species (left), and age of families with at least two representative genera (right) estiamted by TreePL+CASTLES-Pro and TreePL+ConBL. The direction of the arrow is from ConBL-based dates to CASTLES-Pro dates.

**Figure S18:**
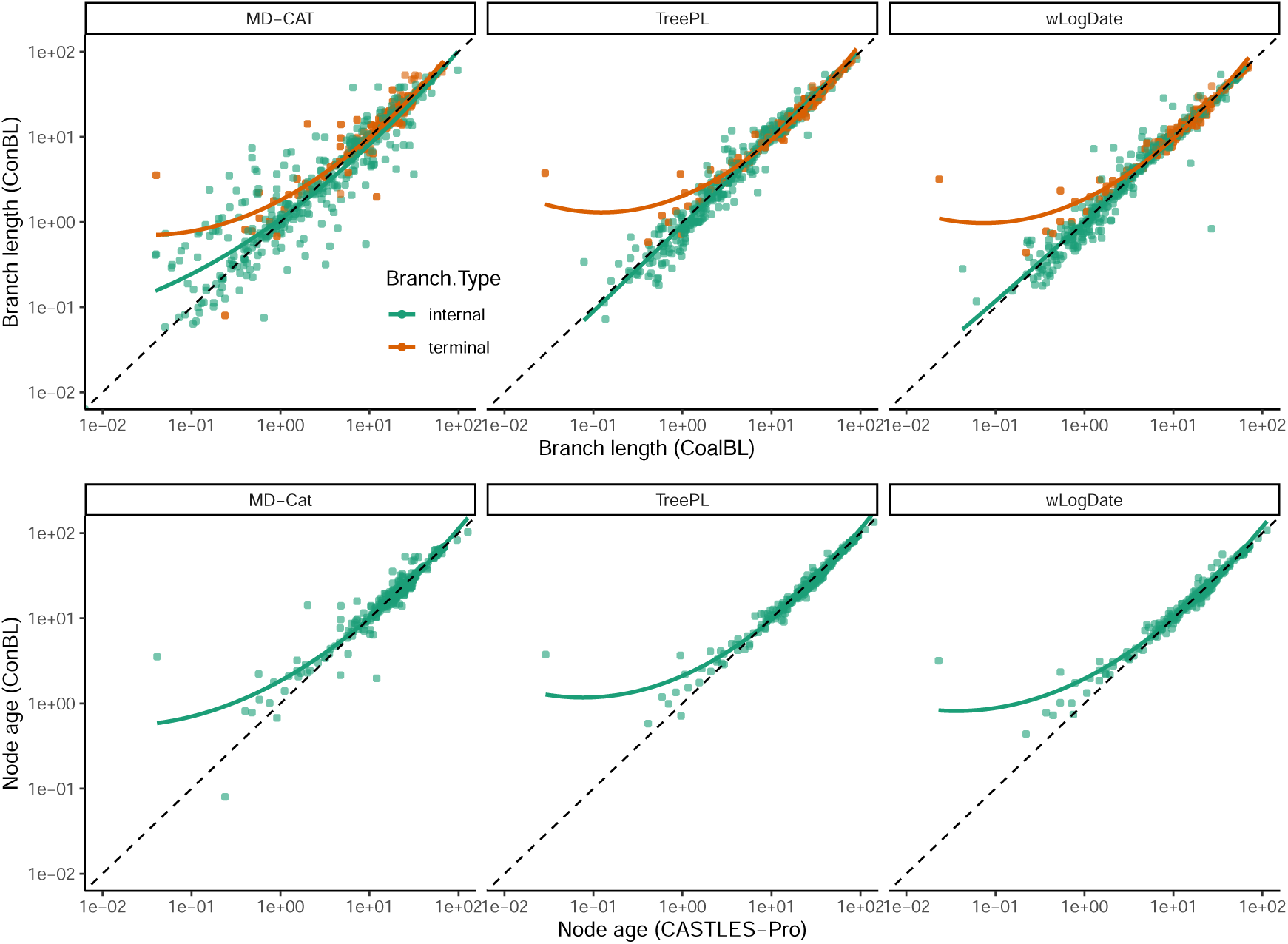
Correlation between time-unit branch lengths (top) and node ages (bottom) of different ML-based dating pipelines using CASTLES-Pro or ConBL on the 363-taxon neoavian phylogeny from Stiller et al. (2024). The tree topology is the main ASTRAL topology from the original study estimated from 63K gene trees. MD-Cat, TreePL, and wLogDate are run with fixed calibration points using median quantiles of calibration densities from the main study. The direction of time in the bottom plot is considered backward so that all extant (terminal) taxa have age 0 and internal nodes have non-zero ages, and so only the age of internal nodes are shown. Both axes are in log scale.

**Figure S19:**
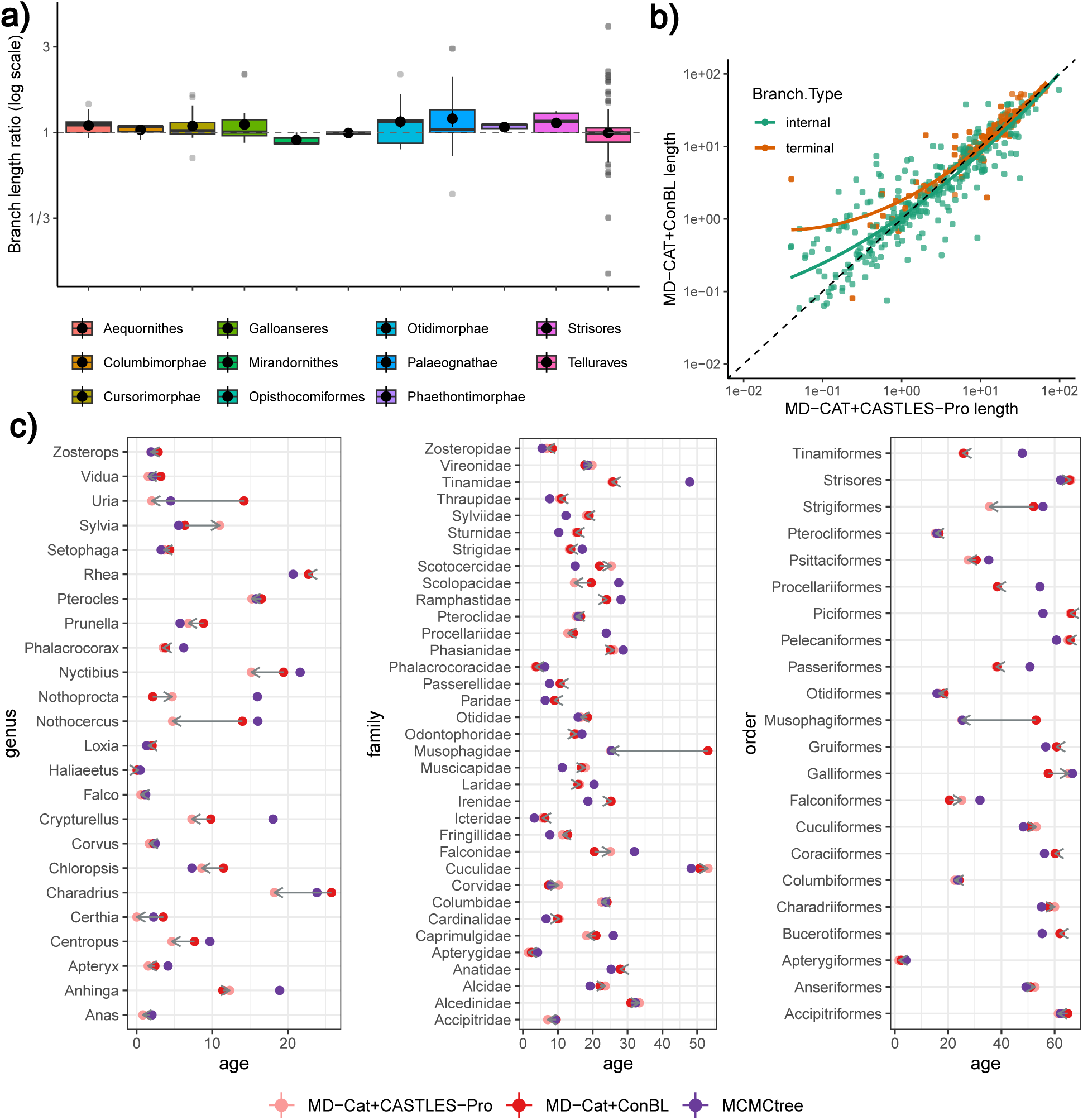
Results of dating the 363-taxon neoavian phylogeny (Stiller et al., 2024) using MD-Cat on an ASTRAL topology from the original study. a) Terminal branch lengths of MD-Cat+ConBL divided by terminal branch lengths of MD-Cat+CASTLES-Pro in log2 scale across 11 higher order clades. b) Correlation between all branch lengths estimated by MD-Cat+ConBL and MD-Cat+CASTLES-Pro. c) Age of the genera that have at least two representative species, and age of families and orders with at least three representative species estimated by MD-Cat+CASTLES-Pro, MD-Cat+ConBL, as well as MCMCtree analysis from the original study. The direction of the arrow is from ConBL dates to CASTLES-Pro dates.

**Figure S20:**
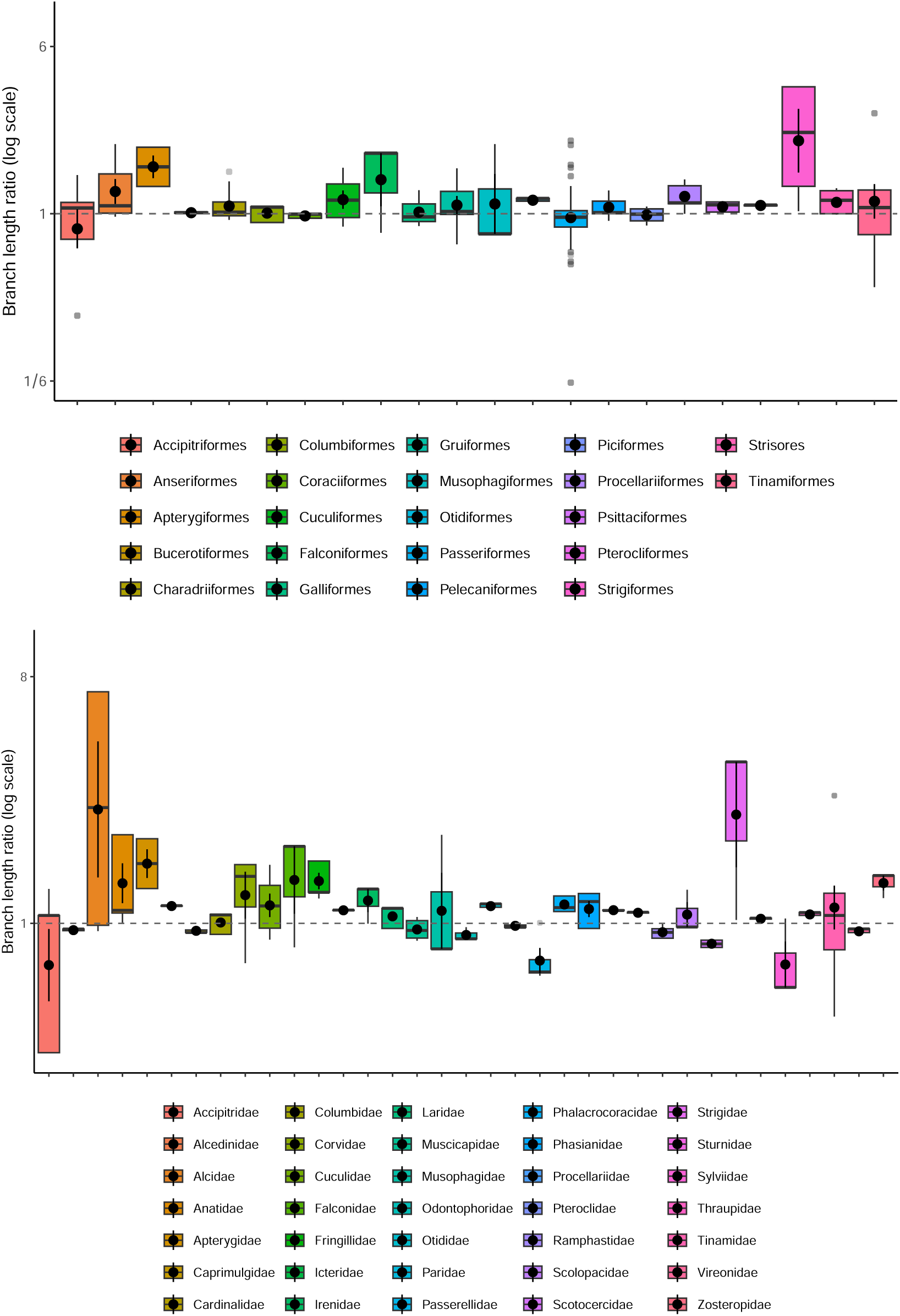
Ratio between terminal branch lengths of trees dated using MD-Cat based on ConBL and CoalBL for the 363-taxon neoavian phylogeny (Stiller et al., 2024) on the ASTRAL topology from the original study. The results show terminal branch lengths of MD-Cat+ConBL divided by terminal branch lengths of MD-Cat+CASTLES-Pro across orders (top) and families (bottom) with more than two representative species in the log2 scale. The panels show the mean (shown with a circle) and standard deviation in addition to boxplots.

**Figure S21:**
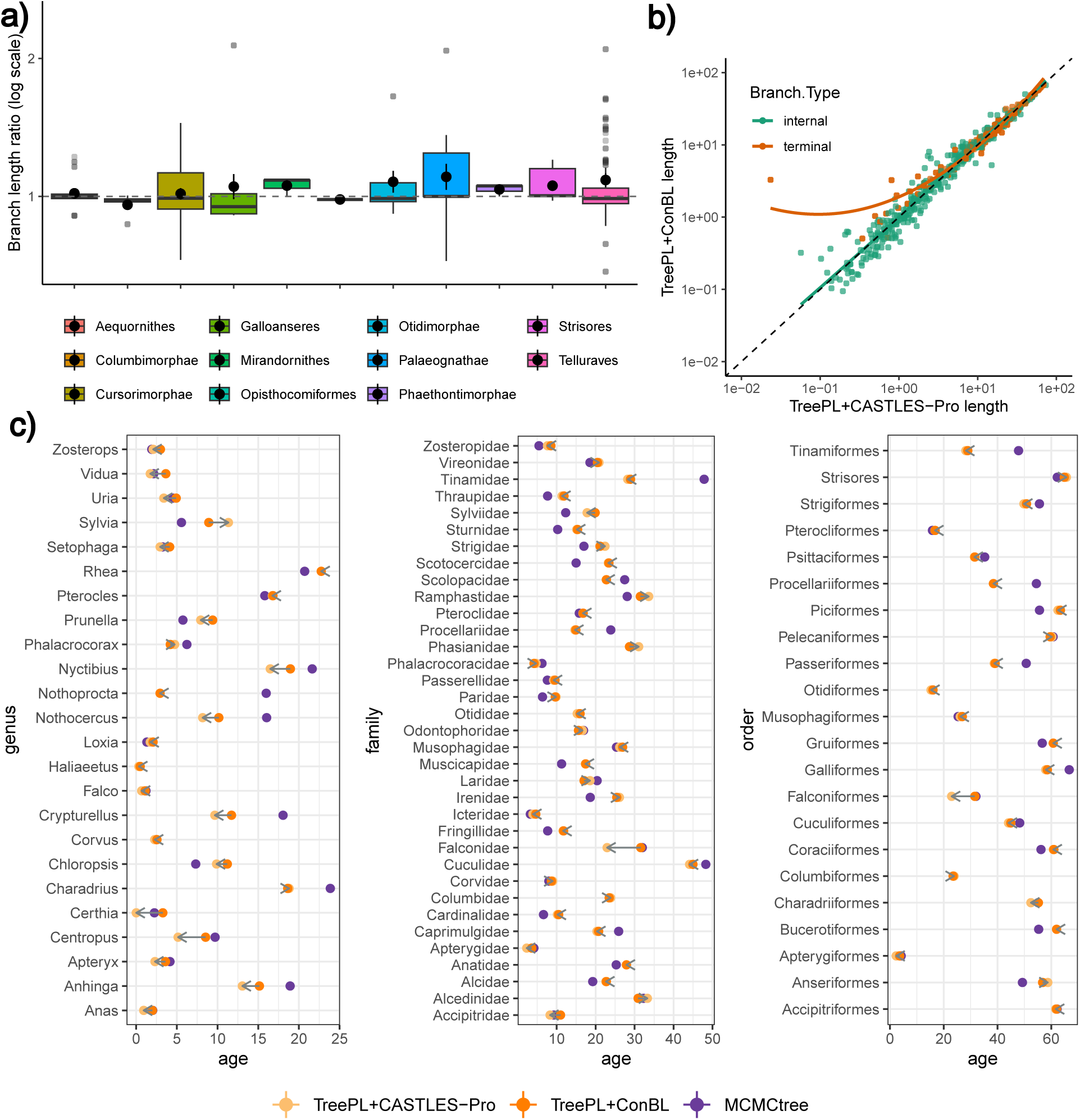
Results of dating the 363-taxon neoavian phylogeny (Stiller et al., 2024) using TreePL on an ASTRAL topology from the original study. a) Terminal branch lengths of TreePL+ConBL divided by terminal branch lengths of TreePL+CASTLES-Pro in log2 scale across 11 higher order clades. b) Correlation between all branch lengths estimated by TreePL+ConBL and TreePL+CASTLES-Pro. c) Age of the genera that have at least two representative species, and age of families and orders with at least three representative species estimated by TreePL+CASTLES-Pro, TreePL+ConBL as well as MCMCtree analysis from the original study (which used a subset of loci used in other analyses, and hence is not directly comparable). The direction of the arrow is from ConBL dates to CASTLES-Pro dates.

**Figure S22:**
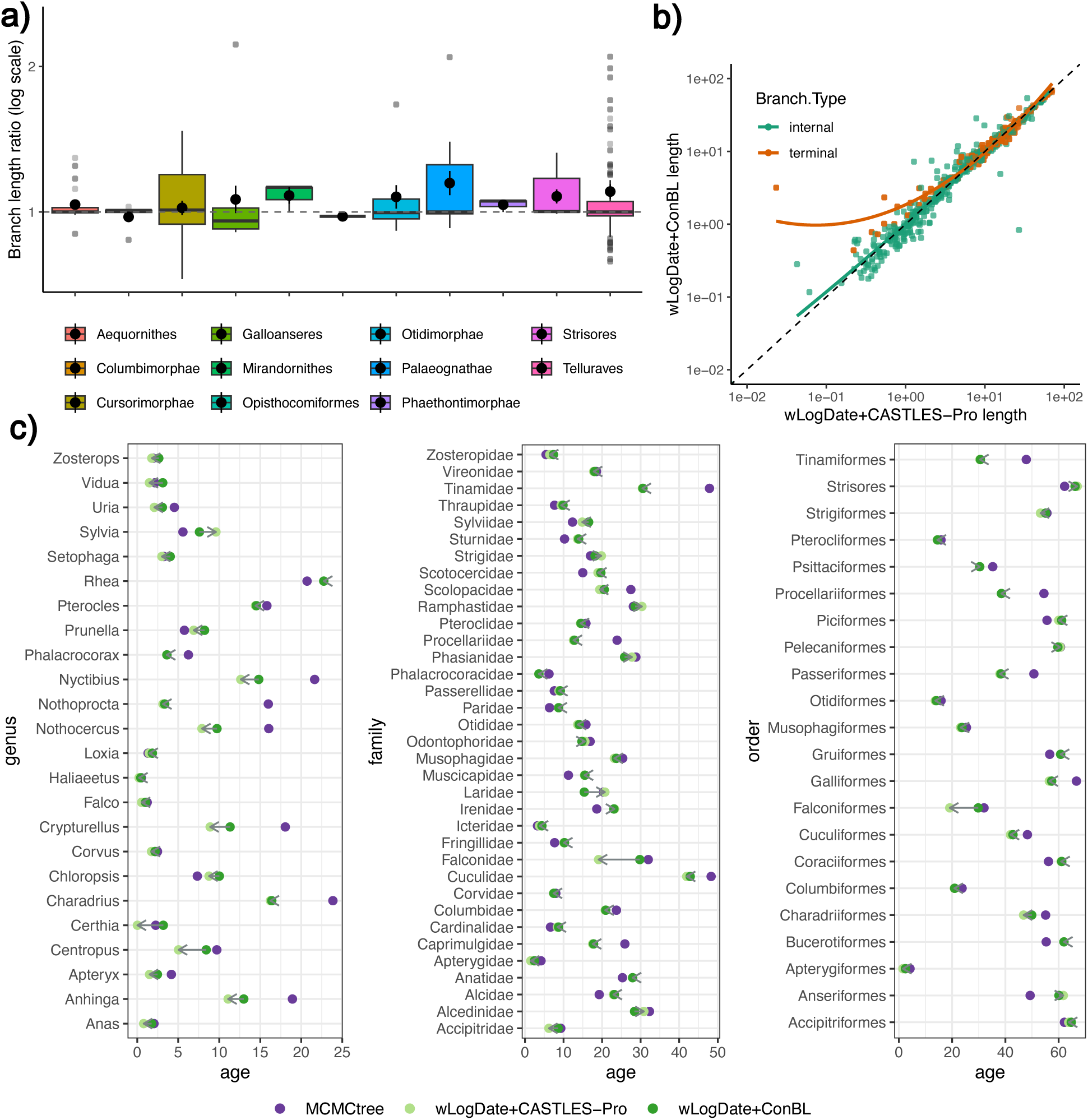
Results of dating the 363-taxon neoavian phylogeny (Stiller et al., 2024) using wLogDate on an ASTRAL topology from the original study. a) Terminal branch lengths of wLogDate+ConBL divided by terminal branch lengths of wLogDate+CASTLES-Pro in log2 scale across 11 higher order clades. b) Correlation between all branch lengths estimated by wLogDate+ConBL and wLogDate+CASTLES-Pro. c) Age of the genera that have at least two representative species, and age of families and orders with at least three representative species estiamted by wLogDate+CASTLES-Pro, wLogDate+ConBL as well as MCMCtree analysis from the original study. The direction of the arrow is from ConBL dates to CASTLES-Pro dates.

**Table S3:**
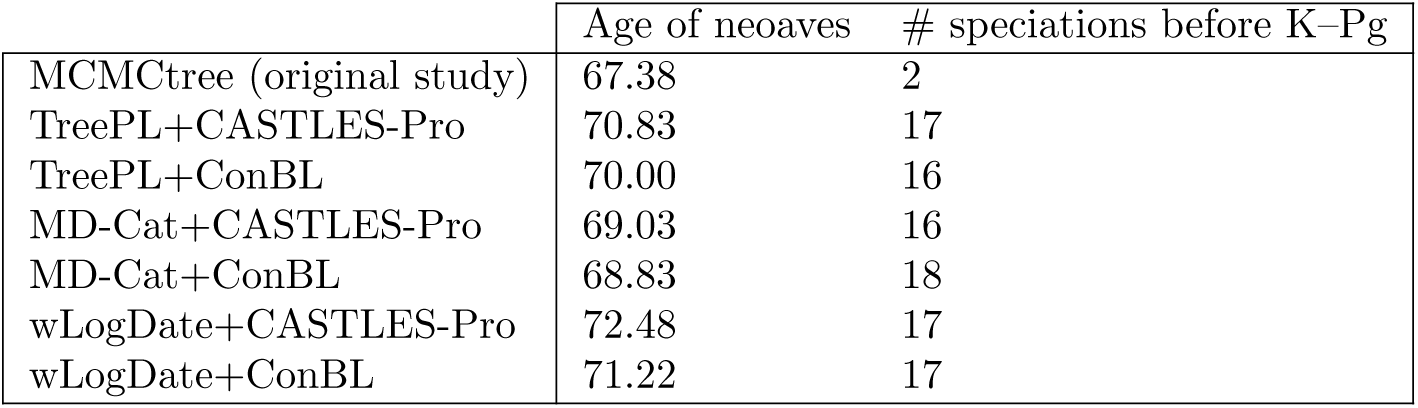
Age of the neoaves as estimated by different dating pipelines (MCMCtree and ML-based dating using CASTLES-Pro and ConBL branch lengths) and the number of speciations before the K–Pg boundary on the 363-taxon dataset of Stiller et al. (2024). Dating is done on an ASTRAL topology from the original study.

|  | Age of neoaves | # speciations before K-Pg |
| --- | --- | --- |
| MCMCtree (original study) | 67.38 | 2 |
| TreePL+CASTLES-Pro | 70.83 | 17 |
| TreePL+ConBL | 70.00 | 16 |
| MD-Cat+CASTLES-Pro | 69.03 | 16 |
| MD-Cat+ConBL | 68.83 | 18 |
| wLogDate+CASTLES-Pro | 72.48 | 17 |
| wLogDate+ConBL | 71.22 | 17 |

## Notes

https://github.com/ytabatabaee/coalescent-based-dating

### Competing Interest Statement

The authors have declared no competing interest.

